# ADSS1 is a suppressed metabolic control node in dystrophic skeletal muscle

**DOI:** 10.64898/2026.07.30.739447

**Authors:** Emma Rybalka, Stephanie Kourakis, Bo Qi, Dean G. Campelj, Ankita Tulangekar, Gretel S. Major, Nitika Kandhari, Amanda Peterson, Christos G. Stathis, Brunda Nijagal, Deanna Deveson-Lucas, Dirk Fischer, Jeffrey J. Brault, Angus Lindsay, Hyung Jun Park, Cara A. Timpani

## Abstract

Duchenne muscular dystrophy is initiated by dystrophin loss but is accompanied by profound metabolic remodelling. Here, we identify suppression of the muscle-enriched adenylosuccinate synthase, ADSS1, and the purine nucleotide cycle in *mdx* muscle. Integrated multi-omic profiling revealed a compensated purine-stress state in which nucleotide salvage and quality control were increased to preserve adenine nucleotide abundance and buffer consequential toxic/disruptive deoxy-/nucleotide production. Purine metabolism was the highest-impact joint pathway with metabolite and transcript hits in *mdx* muscle, and its maintenance program was directionally conserved in human ADSS1 myopathy. Interventions positioned around ADSS1 separated the outputs of this node. Ribose and dimethyl fumarate remodelled upstream or downstream stress programs, whereas only adenylosuccinic acid bypassed ADSS1 to expand the adenine nucleotide pool and remodel CoA/acetyl-CoA metabolism. All three interventions suppressed pro-adipogenic transcription without broadly correcting the lipidome. These findings identify ADSS1 as a regulated metabolic control node that couples purine retention to inflammatory and adipogenic remodelling in dystrophic muscle.

## Introduction

Genetic disease can expose metabolic functions that remain obscured under normal physiological conditions. The identification of adenylosuccinate synthase isozyme 1 myopathy (ADSS1 myopathy) in 2016 revealed that genetic disruption of a single reaction in purine metabolism is sufficient to impair skeletal muscle function, manifesting as progressive weakness, wasting, fatigue and intramuscular fat replacement^1^. ADSS1 expression or activity is also reduced in other neuromuscular disorders not caused by *ADSS1* variants, including Duchenne muscular dystrophy (DMD)^2,3^, amyotrophic lateral sclerosis (ALS)^4,5^ and Huntington’s disease^6^. Because ADSS1 catalyses the conversion of inosine monophosphate (IMP) and aspartate to adenylosuccinate and is strongly enriched in skeletal muscle^7,8^, these observations raise the possibility that reduced ADSS1 activity is part of a broader metabolic response to chronic cellular stress.

Skeletal muscle must accommodate rapid changes in ATP demand while preserving adenine nucleotide (AdN) balance. Adenylate kinase (AK) buffers this demand by catalysing the reversible interconversion of ATP, ADP and AMP. During intense contraction, AMP deaminase 1 (AMPD1) converts AMP to IMP, helping to maintain the AK reaction and ATP:ADP ratio at the cost of transient reduction of the AdN pool. IMP can subsequently be returned to AMP through sequential reactions catalysed by ADSS1 and adenylosuccinate lyase (ADSL) or degraded and lost from the nucleotide pool altogether. Collectively, ADSS1, ADSL and AMPD1 constitute the muscle-enriched purine nucleotide cycle (PNC; Figure 1A). The ADSS1-ADSL reaction sequence also consumes aspartate and releases fumarate, linking AdN metabolism with cellular carbon and nitrogen handling^7^.

**Figure 1.**
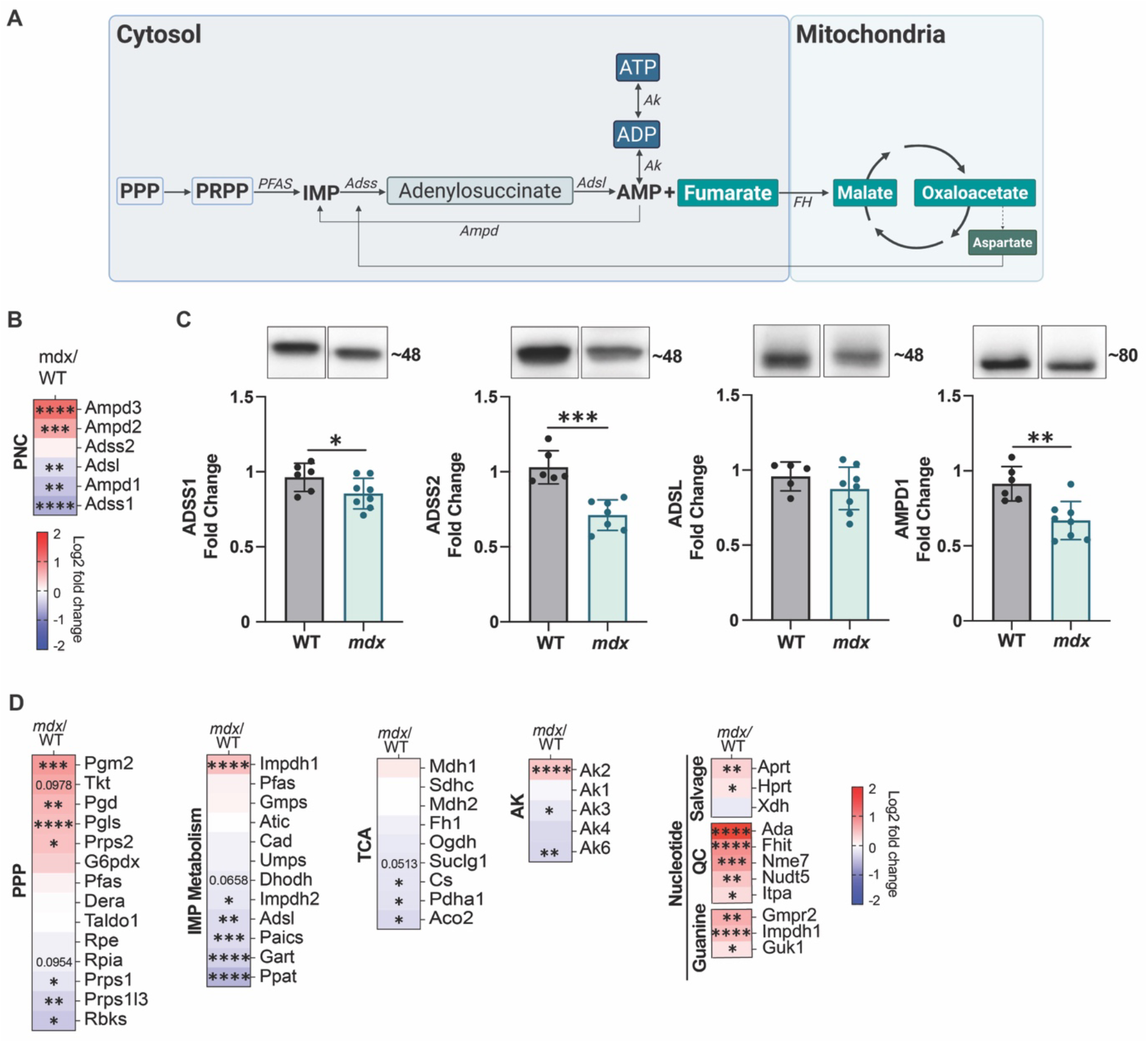
The purine nucleotide cycle is suppressed within a nucleotide-conservation program in dystrophic *mdx* muscle. (A) Schematic of the ADSS1 metabolic node and associated pathways. (B) RNA-sequencing analysis of the purine nucleotide cycle (PNC) genes. (C) Protein abundance of adenylosuccinate synthase 1 (ADSS1), ADSS2, adenylosuccinate lyase (ADSL) and adenosine monophosphate deaminase (AMPD1). Full length blots are shown in Figure S1. (D) Transcriptional changes in pathways associated with pentose phosphate metabolism (PPP), inosine monophosphate (IMP) synthesis, purine salvage and catabolism, adenylate kinase (AK), and the mitochondrial tricarboxylic acid (TCA) cycle. \**p*<0.05, \*\**p*<0.01, \*\*\**p*<0.001, \*\*\*\**p*<0.0001 *mdx* different from wildtype (WT). Statistical trends (*p*=>0.05, <0.1) are labelled with exact *p* values.

The physiological consequences of disrupting different PNC reactions are not equivalent. AMPD-deficiency alleles are common and frequently tolerated^9^, whereas biallelic variants that impair ADSS1 cause a progressive myopathy^1^. ADSL deficiency causes a predominantly neurological disorder because ADSL participates in both AMP synthesis and *de novo* IMP biosynthesis, although muscular dystrophy has also been reported^10^. This asymmetry suggests that the capacity to regenerate AdN’s from IMP may be particularly important for maintaining muscle function during recurrent energetic demand. ADSS1 has also been linked to HDAC3-dependent regulation of metabolic gene expression^11^, raising the possibility that this reaction – or moonlighting functions of the protein – influences cellular adaptation beyond exclusively nucleotide biosynthesis.

DMD is caused by loss of dystrophin and mechanical instability of the sarcolemma^12^, but dystrophic muscle also exhibits altered mitochondrial function, substrate use and energetic regulation^13^. Metabolic abnormalities have additionally been described in dystrophin-deficient muscle stem cells before dystrophin is normally expressed^14^, suggesting that they are not exclusively a consequence of myofibre rupture. The finding that ADSS1 is reduced in DMD muscle^2^, together with the overlap between DMD-related transcriptional pathways and those altered in ADSS1 myopathy^15^, raises a specific question: does ADSS1 suppression form part of the metabolic adaptation of dystrophic muscle, and does it constrain the capacity of muscle to respond to sustained energetic stress?

Here, we combined transcriptomic, metabolomic and lipidomic profiling in an exercise-exacerbated *mdx* mouse model^16^ to define the molecular context of ADSS1 suppression in dystrophic skeletal muscle. We then compared interventions positioned upstream (ribose), across (adenylosuccinic acid; ASA) and downstream ((dimethyl) fumarate; DMF) of the ADSS1 reaction. ADSS1 suppression occurred within a coordinated program of reduced *de novo* nucleotide-biosynthetic machinery, preserved basal AdN abundance and increased salvage and quality-control. Only ASA expanded the AdN pool, demonstrating ADSS1 is pharmacologically tractable. Cross-species integration further identified a conserved purine-maintenance program linking *mdx* and human ADSS1 myopathy muscle.

## Results

### PNC suppression in dystrophic *mdx* muscle accompanies a nucleotide-conservation program

Reduced *ADSS1* expression has been reported in muscles from DMD patients^2^. To determine whether a similar change was present in *mdx* muscle, we quantified PNC gene and protein expression in quadriceps from 8-week-old wildtype (WT) and exercise aggravated *mdx* mice that had previously been characterised both functionally and histopathologically^16^. Transcripts encoding *Adss1*, *Adsl* and *Ampd1*, were lower in *mdx* muscle (Figure 1B). ADSS1 and AMPD1 protein abundance was correspondingly reduced, whereas ADSL protein abundance was maintained despite lower transcript expression (Figure 1C and Figure S1).

We next examined whether paralogous enzymes compensated for suppression of the muscle-enriched PNC. *Ampd2* and *Ampd3*, which are expressed more broadly than *Ampd1* and can be enriched in proliferating or non-myofibre cell populations, were higher in *mdx* muscle (Figure 1B). *Adss2* transcript abundance was unchanged, but ADSS2 protein abundance was lower (Figure 1B, C and Figure S1). Thus, the muscle PNC was suppressed without evidence of compensatory ADSS2 induction at either the transcript or protein level. The increase in *Ampd2* and *Ampd3* may instead reflect isoform switching and/or altered cellular composition in dystrophic muscle.

To place PNC suppression within the wider nucleotide network, we examined transcriptional programs associated with ribose-5-phosphate and phosphoribosyl pyrophosphate (PRPP) production, *de novo* IMP biosynthesis, purine salvage and catabolism, AK and mitochondrial carbon metabolism (Figure 1D). Multiple transcripts encoding *de novo* IMP synthesis and PPP enzymes were lower. In contrast, the purine salvage enzymes *Aprt* and *Hprt* and several nucleotide quality-control genes, including *Ada*, *Itpa*, *Nudt5* and *Fhit* that are responsible for clearing the toxic/disruptive deoxy-/nucleotides deoxy-adenosine, ITP, dITP, ADP-ribose and diadenosine polyphosphates, respectively, were increased. *Prps1* was reduced whereas *Prps2* was increased, indicating isoform-selective remodelling of PRPP synthesis. This pattern is consistent with reduced biosynthetic investment and greater capacity for nucleotide recycling capacity, although neither PRPP abundance nor pathway flux was measured. *Impdh1*, which directs IMP toward guanine nucleotide synthesis, was lower, whereas the proliferation-associated isoform *Impdh2* was higher. Cell purine degradation and excretion enzyme, *Xdh* (xanthine dehydrogenase/oxidoreductase), expression was unchanged.

AK isoforms were also selectively remodelled. The muscle-enriched cytosolic isoform *Ak1* was unchanged, mitochondrial intermembrane space *Ak2* was increased, and mitochondrial matrix *Ak3* and nuclear *Ak6* were reduced (Figure 1D). This pattern suggests compartment-specific adaptation of AdN equilibration rather than uniform induction or suppression of AK capacity. Transcripts encoding citrate synthase (*Cs*), pyruvate dehydrogenase E1α (*Pdh1a*) and mitochondrial aconitase (*Aco2*) were lower, with *Suclg1* trending lower (*p*=0.0513), indicating transcriptional remodelling of pyruvate oxidation and the proximal TCA cycle. These data do not, however, directly establish reduced mitochondrial flux.

Together, these findings place ADSS1 and PNC suppression within a coordinated nucleotide-conservation program in dystrophic muscle. The program combines reduced expression of energetically costly *de novo* biosynthetic machinery with increased expression of salvage and nucleotide quality-control pathways, while retaining selected mechanisms for AdN equilibration.

### Multi-omic profiling identifies compensated purine stress and altered lipid handling in dystrophic muscle

To define the metabolic state associated with ADSS1 suppression, we performed untargeted HILIC metabolomics and lipidomics alongside bulk RNA sequencing of *mdx* quadriceps. Of the 169 detected polar metabolites, 22 differed between *mdx* and WT muscle, with 11 increased and 12 decreased (Figure 2A and Table S1). Kynurenine and anthranilate were increased, consistent with inflammatory activation of tryptophan metabolism. Inosine triphosphate (ITP), a non-canonical purine nucleotide removed by ITPA-dependent quality control, was also increased. O-phosphoethanolamine (PEA) and glycerophosphocholine (GPC) were higher, indicating altered phospholipid metabolism or membrane turnover. Metabolites reduced in *mdx* muscle included adenine, NADH, carnosine, glycine and other amino acids that contribute to nitrogen and one-carbon metabolism. Pathway analysis highlighted enrichment of ammonia recycling, purine metabolism and membrane-associated pathways (Figure 2B and Table S2).

**Figure 2.**
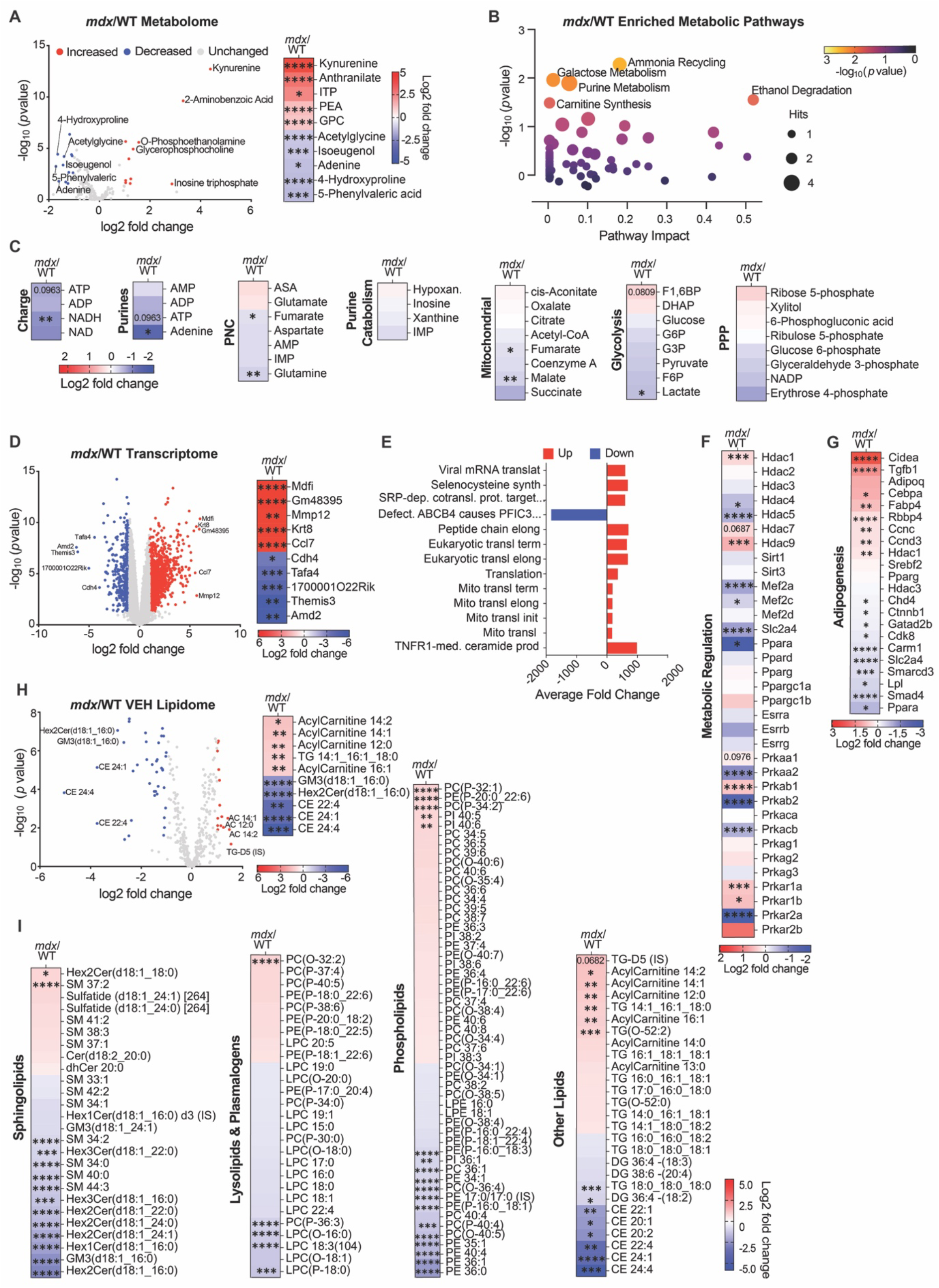
Multi-omic profiling identifies compensated purine stress and altered lipid handling in dystrophic muscle. (A) Volcano plot and heatmap of the five most increased and decreased polar metabolites in *mdx* muscle relative to wildtype (WT). (B) Metabolite-set enrichment analysis. (C) Metabolites associated with charge, purine, purine nucleotide cycle (PNC), tricarboxylic acid (TCA) cycle, glycolytic and pentose phosphate (PPP) pathways. (D) Volcano plot and heatmap of the most strongly altered transcripts. (E) Reactome pathway enrichment. (F) Transcripts encoding selected regulators of metabolic signalling. (G) Adipogenesis- and lipid-regulatory transcripts. (H) Volcano plot of the lipidome. (I) Lipid species grouped by class. Complete significantly regulated metabolite, pathway and lipid datasets, as well as the top fifty significantly regulated transcripts are provided in Tables S1– S4. \**p*<0.05, \*\**p*<0.01, \*\*\**p*<0.001 and \*\*\*\**p*<0.0001 *mdx* different from WT. Statistical trends (*p*=>0.05, <0.1) are labelled with exact *p* values.

ATP showed a non-significant downward trend (*p*=0.0963), but ADP and AMP were not significantly reduced between *mdx* and WT muscle (Figure 2C). NADH but not NAD+ was also significantly reduced. These findings do not indicate overt loss of basal AdN abundance or energetic collapse at the sampled time point. Instead, reduced adenine together with increased ITP and induction of salvage and quality-control genes indicates pressure on purine maintenance despite preservation of the AdN abundance. Among other PNC- and central-carbon-associated metabolites, adenine, fumarate, malate, glutamine and lactate were lower, whereas adenylosuccinate, aspartate and IMP were unchanged (Figure 2C).

The most strongly altered transcripts primarily reflected chronic injury and cellular remodelling, e.g., increased *Mmp12*, *Krf8* and *Ccl7* (Figure 2D and Table S3). Pathway enrichment identified cytosolic and mitochondrial translational pathways among the most affected pathways (Figure 2E). In the absence of coordinated induction of mitochondrial biogenesis or respiratory-complex assembly, this signature is more consistent with remodelling of protein synthesis and mitochondrial stress responses than with increased mitochondrial capacity. TNFR1-mediated ceramide production was among the most strongly upregulated pathways, linking metabolic phenotype to inflammatory and sphingolipid signalling.

We next examined transcriptional regulators of muscle metabolism (Figure 2F). *Mef2a* and *Mef2c* were reduced by >1.2 fold, together with extensive but non-uniform changes in histone deacetylase expression. The AMP-activated protein kinase (AMPK) complex was also remodelled: *Prkab1* was increased, whereas *Prkaa2* and *Prkab2* were reduced. Within the cAMP-dependent protein kinase A (PKA) system, *Prkacb* and *Prkar2a* were reduced, and *Prkar1a* was increased. These data indicate reorganisation of metabolic signalling machinery but not necessarily altered AMPK or PKA activity, which depends on each of nucleotide concentration, phosphorylation, subcellular localisation and holoenzyme composition rather than transcript abundance alone.

Because intramuscular fat replacement is prominent in human DMD, we examined lipid-regulatory programs to assess whether early molecular signatures of adipogenic remodelling were present, despite the lack of widespread histologically evident adiposis in contralateral quadriceps of these mice^16^. *Mdx* muscle nevertheless exhibited a strong adipogenesis-associated transcriptional signature (Figure 2G). *Cidea*, *Tgfb1*, *Cebpa* and *Fabp4* were increased, whereas *Ppara*, *Smad4*, and *Lpl*, and several chromatin-regulatory genes (*Smarcd3*, *Carm1*, *Cdk8*, *Gatad2b*, *Ctnnb1*, *Chd4*), were reduced. Lipidomics showed accumulation of multiple medium-chain acylcarnitines and triacylglycerols and depletion of several membrane-associated lipid classes (Figure 2J and K and Table S4). This pattern indicates altered fatty-acid handling and activation of a pro-adipogenic or stromal remodelling program albeit without histological adipocyte accumulation.

Despite dystrophic muscle maintaining its basal AdN pool, it expresses molecular evidence of purine stress, altered central-carbon metabolism, inflammatory activation and lipid-handling defects. Preservation of AdN abundance in this context is consistent with effective compensation, including increased nucleotide salvage capacity, albeit decreased adenine suggests compensation stress.

### ASA, DMF and ribose produce distinct transcriptional responses around the ADSS1 node

To interrogate different components of ADSS1-centred metabolism, *mdx* mice were treated with interventions positioned around the ADSS1 reaction sequence (Figure 3A). Ribose was used to increase upstream pentose availability for ribose-5-phosphate, PRPP and ultimately, IMP production as a substrate for ADSS1; ASA supplied the product of the ADSS1 reaction; and DMF was used to engage fumarate- and redox-responsive pathways without supplying an adenine nucleotide precursor. Ribose produced the largest transcriptional response relative to vehicle-treated *mdx* muscle, whereas ASA and DMF produced smaller, partly overlapping signatures (Figure 3B and C).

**Figure 3.**
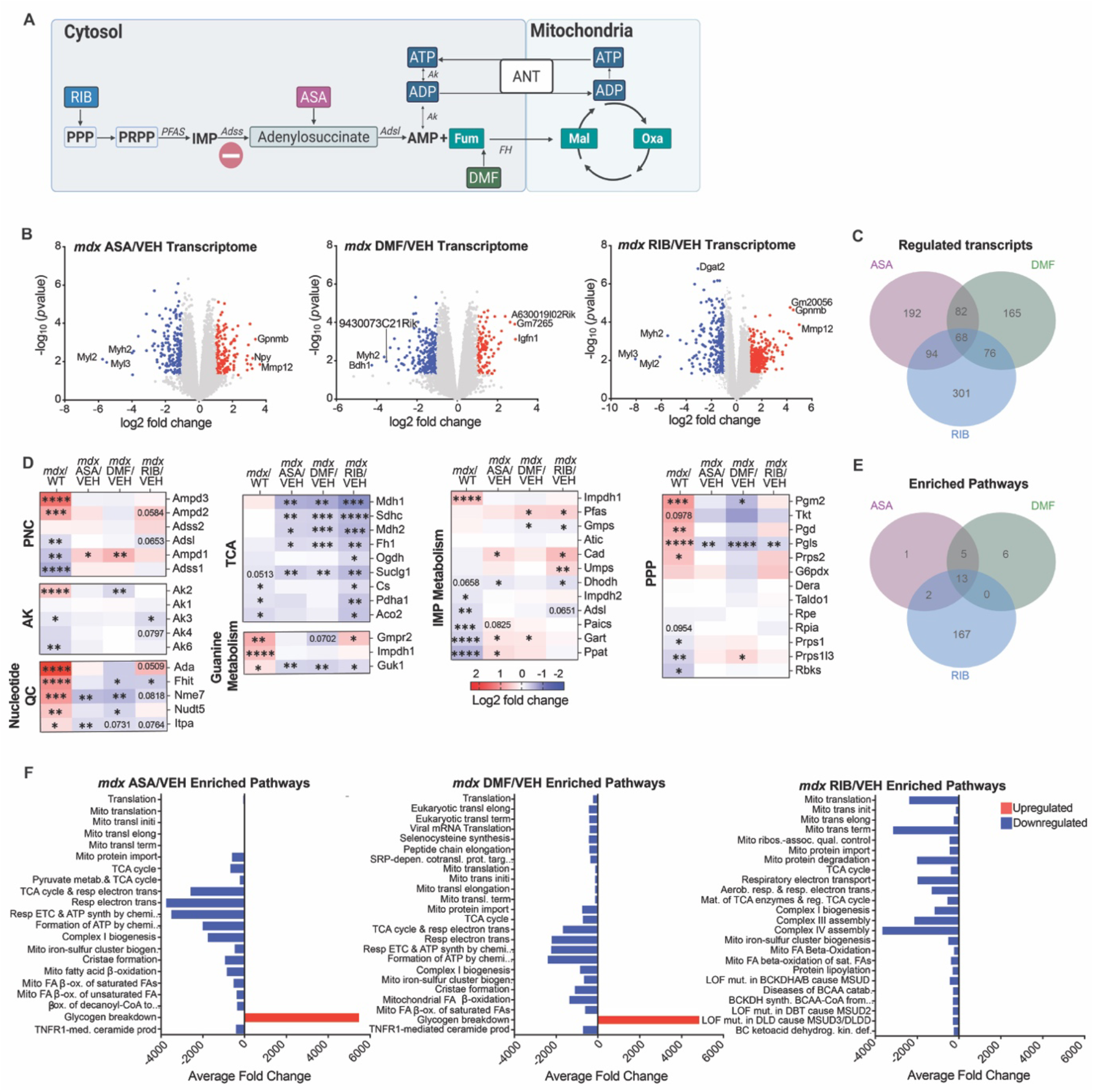
ASA, DMF and ribose differentially remodel transcriptional programs associated with the ADSS1 node. (A) Schematic showing the metabolic positions of adenylosuccinic acid (ASA), dimethyl fumarate (DMF) and ribose (RIB). (B) Volcano plots of treatment-induced transcriptional changes relative to vehicle-treated *mdx* muscle. (C) Overlap among differentially expressed transcripts. (D) Selected nucleotide- and central-carbon-associated transcripts. (E) Overlap among enriched pathways. (F) Comparative pathway enrichment. Enriched pathways based on number of genes mapped is provided in Figure S2. The top fifty up and down regulated transcripts are provided in Table S3. \**p*<0.05, \*\**p*<0.01, \*\*\**p*<0.001 and \*\*\*\**p*<0.0001 treatment different from *mdx* vehicle. Statistical trends (*p*=>0.05, <0.1) are labelled with exact *p* values.

All three interventions altered expression of nucleotide-associated genes, but no single response completely restored the untreated WT program (Figure 3D). Nucleotide quality control gene upregulation in mdx muscle was mostly normalised by the interventions – ASA targeted *Nme7* and *Itpa*, DMF targeted *Nme7* and *Nudt5*, and ribose targeted *Fhit*. Similarly, TCA cycle enzymes were decreased entirely by ribose, and partially (55% of them) by ASA and DMF. ASA and ribose increased *Cad,* which encodes the multifunctional enzyme catalysing the first steps of *de novo* pyrimidine synthesis, and reduced *Dhodh*, which couples mitochondrial electron transfer to pyrimidine synthesis. All interventions reduced *Guk1* and ribose and DMF reduced *Gmps*, indicating remodelling of the IMP-to-GMP branch. All interventions reduced *Pgls* indicating PPP remodelling rather than suppression or restoration of pathway activity. DMF increased *Prps1l3*, whereas ASA and DMF increased *Ampd1*, indicating intervention-specific effects on PRPP-synthesis and PNC gene expression.

Pathway enrichment further distinguished the three treatments (Figure 3E,F). Ribose predominantly affected mitochondrial organisation and metabolic-adaptation pathways, consistent with its broad transcriptional effect. ASA and DMF showed greater overlap in pathways associated with mitochondrial translation and stress responses. Each intervention also reduced selected TCA-cycle transcriptional pathways that were altered in untreated *mdx* muscle.

### ASA expands AdN availability and remodels acetyl-CoA-associated metabolism

We next determined whether the distinct transcriptional responses were accompanied by changes in metabolite abundance (Figure 4A–D). ASA produced the broadest metabolomic response, altering 23 metabolites at the prespecified threshold (|log2FC| ≥ 1.0, *p*< 0.05) and enriching 43 metabolic pathways (Figure 4A-C and Table S5). DMF and ribose altered 4 and 17 metabolites and enriched three and one metabolic pathways, respectively (Figure 4A–C and Tables S6 and S7).

**Figure 4.**
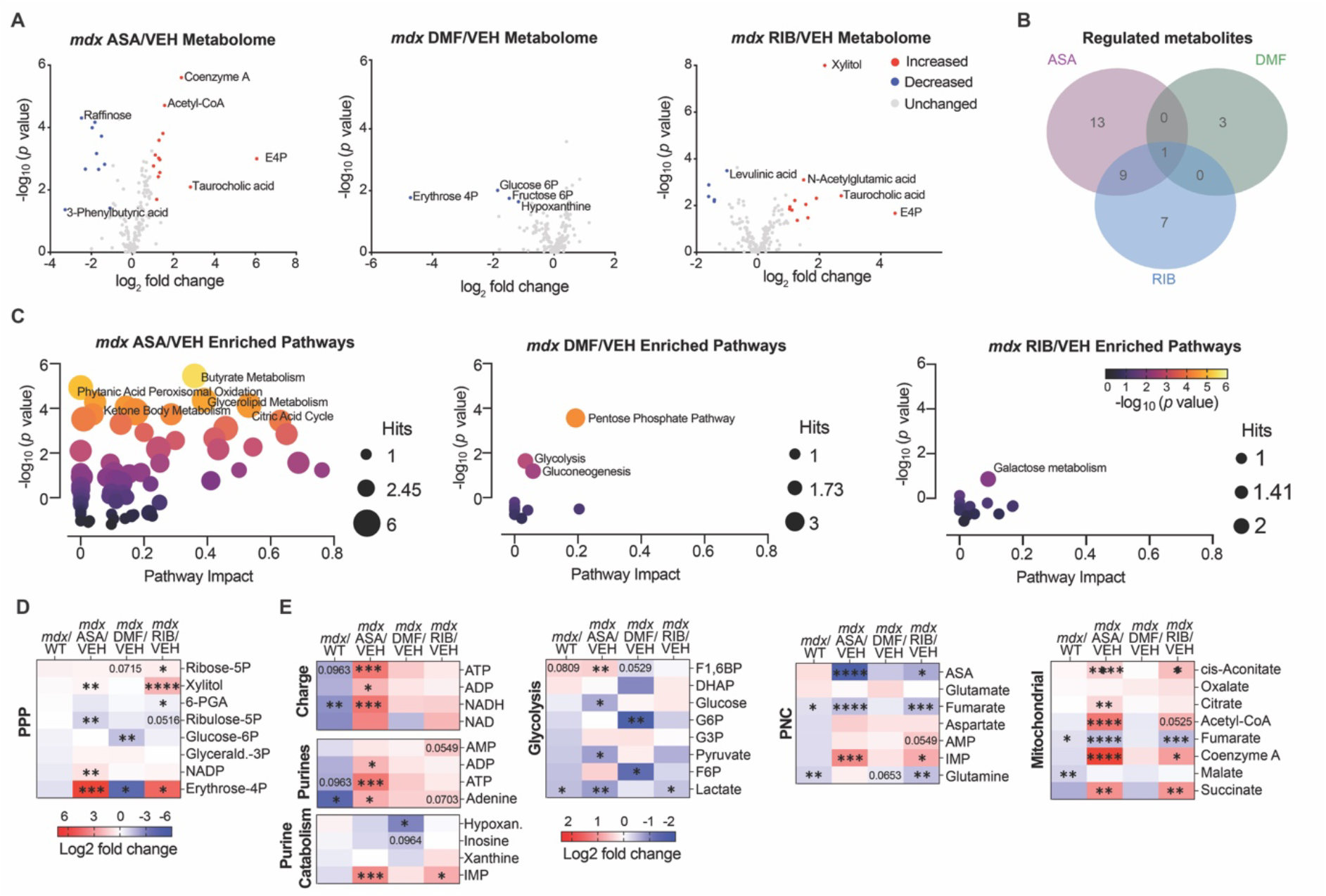
ASA increases adenine nucleotide availability and remodels acetyl-CoA-associated metabolism in dystrophic muscle. (A) Volcano plots of metabolites altered by adenylosuccinic acid (ASA), dimethyl fumarate (DMF) and ribose (RIB) relative to vehicle-treated *mdx* muscle. (B) Overlap among treatment-responsive metabolites. (C) Metabolite-set pathway enrichment. (D-E) Selected metabolites associated with purine, pentose-phosphate, CoA/acetyl-CoA and central-carbon metabolism. Complete significantly different treatment-responsive metabolite and pathway datasets are provided in Tables S1 and S5-S7. \**p*<0.05, \*\**p*<0.01, \*\*\**p*<0.001 and \*\*\*\**p*<0.0001 treatment versus vehicle-treated *mdx* muscle. Statistical trends (*p*=>0.05, <0.1) are labelled with exact *p* values.

Erythrose-4-phosphate (E4P) increased by ASA and ribose and decreased with DMF (Figure 4D and Table S1), demonstrating divergent effects of the interventions on pentose- carbon metabolism. E4P is a low-abundance intermediate of the non-oxidative PPP whose steady-state concentration is influenced by reversible transketolase and transaldolase reactions. Taurocholic acid was also increased by ASA and ribose (Table S1), although the physiological basis of this response in skeletal muscle remains unresolved.

ASA expanded AdN abundance, including increased ATP and ADP (and adenine) (Figure 4E). NADH (but not NAD+) was also increased. DMF and ribose did not reproduce these effects. However, DMF did reduce hypoxanthine, linking downstream fumarate-responsive metabolism to terminal purine handling. ASA also increased coenzyme A (CoA-) and acetyl-CoA-associated metabolites and enriched metabolite sets related to ketone-body, butanoate, glycerolipid and TCA-cycle metabolism (Figure 4E and Table S5). These changes associate greater AdN availability with broader remodelling of acetyl-CoA-centred metabolism. While ribose produced extensive transcriptional remodelling it did not significantly increase AdN abundance (AMP and adenine trended), suggesting that greater pentose availability alone is insufficient to reproduce the ASA-associated metabolic state when applied to suppressed ADSS1. DMF altered selected nucleotide- and stress-associated pathways, and lowered specifically glycolysis intermediates, but likewise did not increase AdN abundance.

Together, these data identify a metabolomic response specific to ASA that includes greater AdN availability and coordinated changes in CoA-/acetyl-CoA-associated metabolism demonstrative of ADSS1-mediated coordination of cytosolic and mitochondrial AdN metabolism.

### ADSS1-centred interventions suppress pro-adipogenic transcription without broadly remodelling the lipidome

ASA, DMF and ribose each strongly reduced *Cidea* and *Fabp4* expression (Figure 5A), demonstrating convergence on lipid-regulatory transcription despite their distinct effects on AdN abundance. This shared response argues against adenylate expansion as the sole mechanism and instead implicates signalling features common to multiple points around the ADSS1 node. ASA reduced *Hdac1*, with a similar trend following ribose treatment (*p*=0.0937).

**Figure 5.**
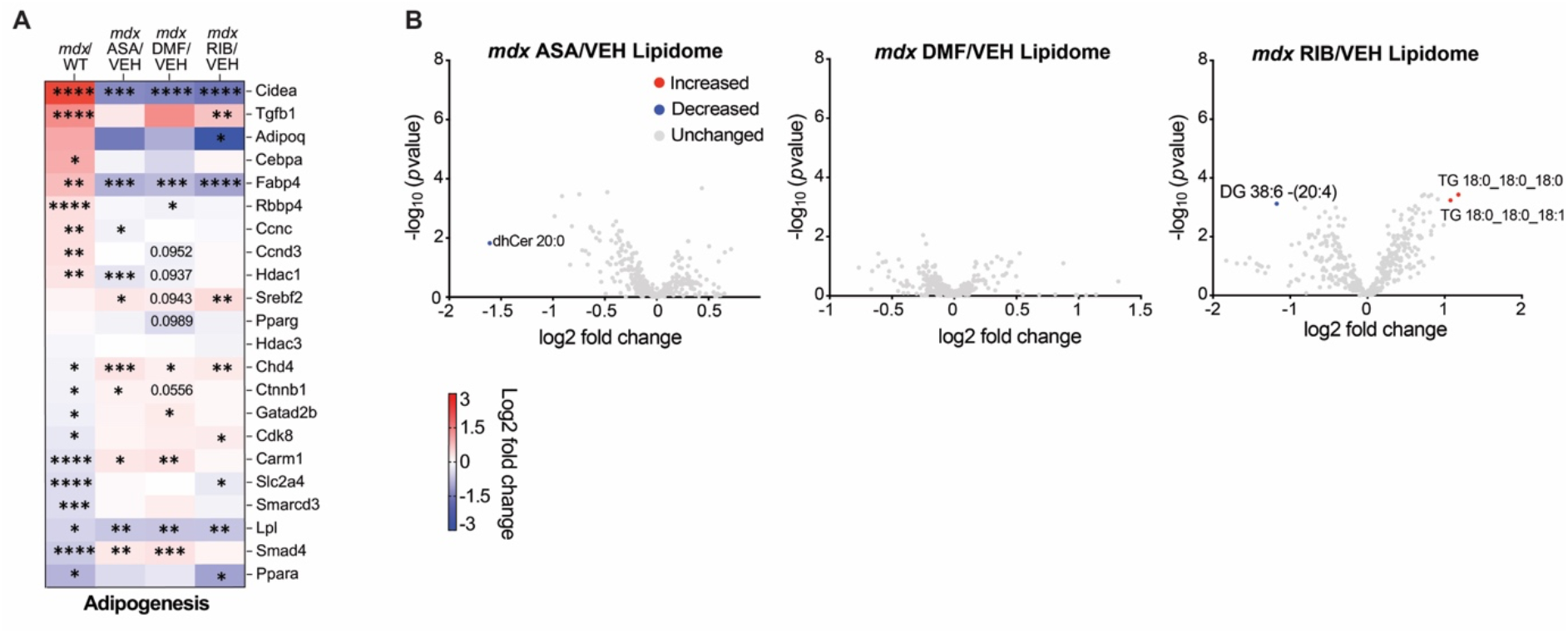
ADSS1-centred interventions suppress pro-adipogenic transcription without broad lipidome remodelling. (A) Adipogenesis- and lipid-regulatory transcripts following adenylosuccinic acid (ASA), dimethyl fumarate (DMF) and ribose (RIB) treatment relative to vehicle-treated *mdx* muscle. (B) Treatment-responsive lipid species. Complete significantly different lipid species are provided in S4. \**p*<0.05, \*\**p*<0.01, \*\*\**p*<0.001 and \*\*\*\**p*<0.0001 treatment versus vehicle-treated *mdx* muscle. Statistical trends (*p*=>0.05, <0.1) are labelled with exact *p* values.

Despite these transcriptional effects, none of the interventions broadly altered the bulk muscle lipidome (Figure 5). ASA selectively reduced dihydroceramide 20:0, whereas ribose increased one and reduced two triacylglycerol species. The limited lipidomic response may reflect slower turnover of established lipid pools, cell-type dilution in whole muscle, or separation between lipid-regulatory transcription and net lipid abundance. Indeed, the widespread fat replacement of muscle tissue in DMD is not recapitulated in murine models of the disease in standard laboratory settings.

### Purine metabolism is the most enriched integrated metabo-transcriptional pathway and anchors a conserved nucleotide maintenance program

A pathway was considered enriched if it contained at least one metabolite and one transcript hit and had an impact score >0.2, with higher impact scores indicating that these alterations occupy more influential positions within the pathway network^17^. Integrated metabolomic-transcriptomic analysis identified *purine metabolism* as the highest-impact pathway containing both metabolite and transcript hits in *mdx* muscle (15 features: 2 metabolites, 13 transcripts), followed by *glycine, serine and threonine metabolism* (6 hits: 1 metabolite, 5 transcripts), *pyrimidine metabolism* (9 hits: 1 metabolite, 8 transcripts), *arginine and proline metabolism* (13 hits: 2 metabolites, 11 transcripts) and *glycerophospholipid metabolism* (12 hits: 2 metabolites, 10 transcripts; Figure 6A and Table S8). ASA altered joint features within *glycerolipid* (4 hits: 2 metabolites, 2 transcripts), *purine* (5 hits: 3 metabolites, 2 transcripts) and *pyrimidine* (4 hits: 1 metabolite, 3 transcripts) metabolism (Figure 6B and Table S9), whereas DMF altered joint features within *purine metabolism* only (6 hits: 1 metabolite, 5 transcripts; Figure 6C and Table S10). Ribose did not meet the prespecified combined-pathway threshold for *purine metabolism* (Figure 6D and Table S11). The interventions therefore engaged the ADSS1-centred network differently, with ASA producing the broadest integrated response.

**Figure 6.**
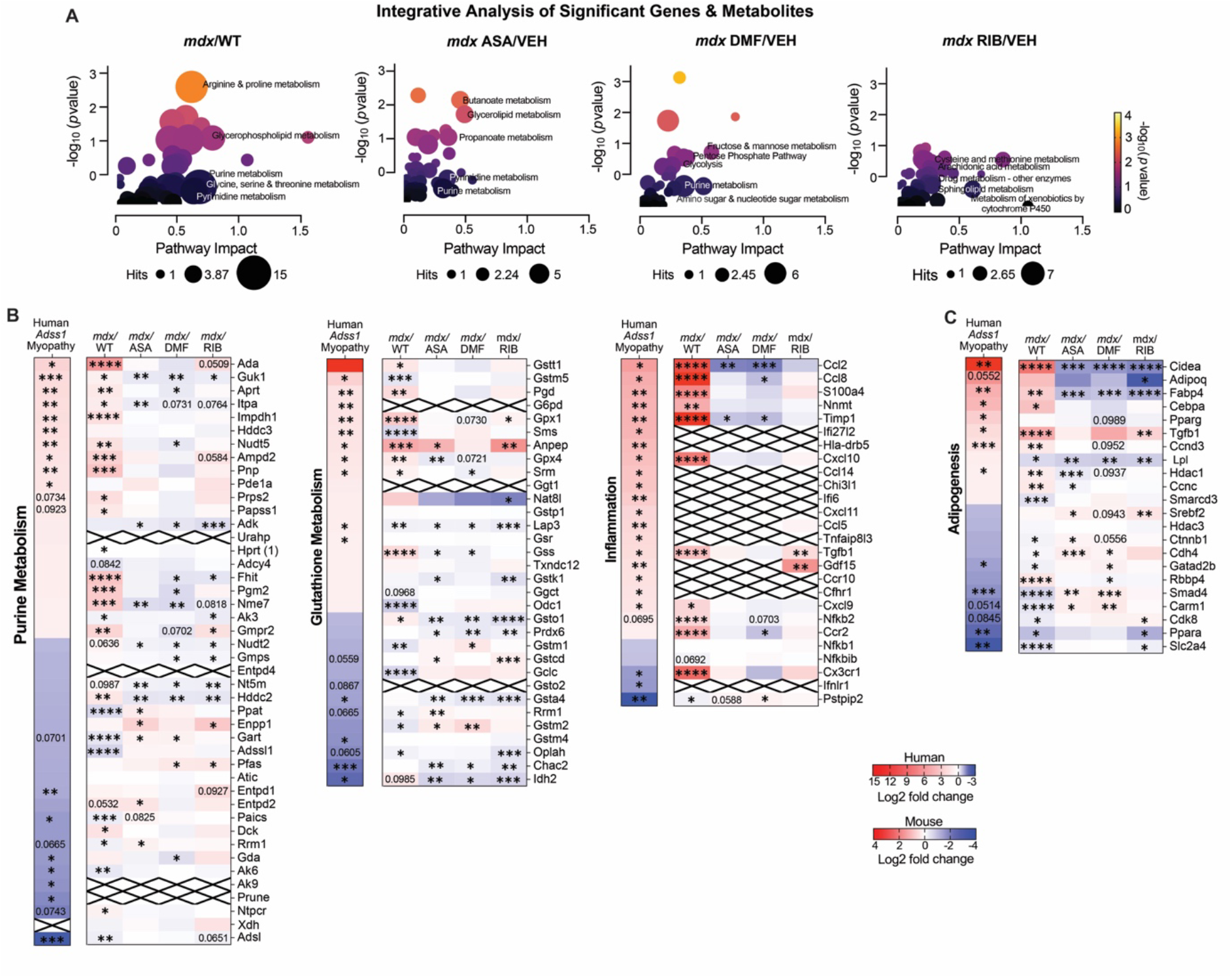
Purine metabolism is the most enriched integrated metabo-transcriptional pathway and anchors a conserved nucleotide maintenance program. (A) Joint metabolomic–transcriptomic pathway analysis of *mdx* versus wildtype (WT) muscle and of adenylosuccinic acid (ASA)-, dimethyl fumarate (DMF)- and ribose (RIB)-treated versus vehicle-treated *mdx* muscle. Displayed pathways met the prespecified impact-score threshold and contained at least one metabolite and one transcript. (B) Cross-species comparison of purine, glutathione, inflammatory and (C) adipogenesis-associated transcriptional modules in *mdx* and human ADSS1-myopathy muscle, together with treatment responses in *mdx* muscle. Integrated pathways datasets are provided in Tables S8–S11. Crosses denote features not detected in a dataset. \**p*<0.05, \*\**p*<0.01, \*\*\**p*<0.001 and \*\*\*\**p*<0.0001 versus the relevant control. Statistical trends (*p*=>0.05, <0.1) are labelled with exact *p* values.

To compare the dystrophic signature with primary ADSS1 deficiency given the overlap of several distinct clinical and histological features, we mapped the *mdx* data to a published transcriptome of skeletal muscle from patients with ADSS1 myopathy^15^. Four metabolic pathways were altered in both *datasets: purine; glycine, serine and threonine; glycerophospholipid; and glutathione* metabolism (Figure S3). Within the purine module, ∼73% of shared differentially expressed genes changed in the same direction across species. Genes with increased transcription were associated with nucleotide salvage or quality control, including *ADA*, *GUK1*, *APRT*, *ITPA*, *IMPDH1*, *NUDT5*, *PRPS2*, *FHIT* and *GMPR2*. In contrast, expression of genes involved in *de novo* nucleotide synthesis were reduced. This conserved pattern supports a shared shift in nucleotide-maintenance capacity, despite the distinct initiating lesions and substantial differences between human ADSS1 myopathy and exercised *mdx* mouse muscle.

We next asked whether interventions opposed components of this shared signature. ASA altered several upstream PPP/*de novo* purine-associated and nucleotide-stress genes, including *Ppat*, *Gart*, *Rrm1, Guk1*, *Itpa*, *Nme7* and *Paics* (trend). DMF more strongly affected a set enriched for salvage-, quality control- and redox-associated genes, including *Aprt*, *Nudt5*, *Fhit*, *Nme7*, *Pgm2* and *Pfas* (Figure 6B). Ribose affected only a few genes significantly and these were genes also modulated by either ASA, DMF or both, e.g., *Guk1 Adk*, *Fhit*. The interventions therefore engaged both shared and distinct components of the conserved program.

Glutathione-, inflammation- and adipogenesis-associated modules also showed substantial directional overlap between *mdx* and human ADSS1 myopathy muscle (Figure 6B and C and Figure S3). These similarities show that primary ADSS1 deficiency and dystrophin-deficient muscle converge on selected stress and tissue-remodelling programs. ASA and DMF opposed induction of *Ccl2* and *Timp1*, and DMF also tempered *Ccl8* (Figure 6B). All three interventions reduced the adipogenic genes, *Cidea* and *Fabp4*, whereas only ASA and DMF tempered lipid metabolism genes *Smad4* and *Carm1*. These cross-species similarities extend the shared purine-maintenance state to selected inflammatory and tissue-remodelling programs.

## Discussion

Our findings reveal that dystrophic muscle did not exhibit resting energetic collapse in the context of ADSS1 suppression. Instead, broad induction of nucleotide salvage and quality control indicate that the AdN pool is dynamically maintained within a reorganised purine network. Purine metabolism was the highest-impact integrated pathway containing both metabolite and transcript hits in *mdx* muscle and provided the strongest metabolic convergence with human ADSS1 myopathy. We used distinct interventions to resolve the functional organisation of this network. Increasing pentose supply or supplying a downstream fumarate analogue remodelled select transcriptional programs, whereas only direct bypass of ADSS1 expanded the AdN pool. These findings identify ADSS1 as a regulated metabolic control node whose suppression changes how dystrophic muscle retains purines, accommodates metabolic demand and coordinates inflammatory and adipogenic remodelling.

### ADSS1 is a regulated metabolic control node

Evidence across experimental systems indicates that residual ADSS activity determines the balance between purine compensation and physiological dysfunction. The approximately 15% reduction of ADSS1 protein in *mdx* muscle was accompanied by adenine depletion and increased salvage, but basal AdN abundance remained intact. A greater 38%–62% reduction of the single *adss-1* isoform in *C. elegans* produces a related metabolic signature – including lower adenine and adenylosuccinate, nominally lower AMP and higher GMP – but with appreciable neuromuscular and growth impairment^18^. In AU-100 lymphoma cells, approximately 80% ADSS deficiency induces *de novo* purine synthesis and more than four-fold IMP accumulation^19^. ATP remains near normal, but GTP increases and excess purine is released predominantly as inosine. Progressive loss of ADSS capacity therefore shifts metabolism from salvage-supported adenylate maintenance towards imbalanced IMP partitioning and purine loss.

Complete loss of ADSS1 reveals the limit of salvage-oriented metabolic remodelling though. Deletion of *adss-1* in *C. elegans* profoundly compromises growth, fertility and neuromuscular function^18^, while loss of yeast ADE12 causes IMP-derived metabolite accumulation, depletion of ATP, pyrimidine nucleotides and dNTPs, G1 arrest and poor survival under adenine limitation^20^. Adenine restores yeast growth, whereas restricting upstream IMP synthesis reduces toxicity, showing that increasing *de novo* synthesis becomes counterproductive when the ADSS bottleneck cannot be relieved and stimulating nucleotide salvage supportive. Mammals retain ADSS2, and its continued expression probably contributes to the relatively mild phenotype of ADSS1-null mice^21^. Dystrophic *mdx* muscle apparently lacks- this predicted reserve since ADSS2 protein was reduced by approximately 30%, twice the reduction in ADSS1.

The conserved salvage response connects graded ADSS1 deficiency models with human disease. *Aprt* and *Hprt* were increased and free adenine was depleted in *mdx* muscle, while approximately 73% of shared purine-pathway genes changed in the same direction in human ADSS1 myopathy muscle. This convergence argues that salvage is recruited in response to restricted ADSS capacity rather than driving the disturbance itself. APRT bypasses ADSS1 by converting adenine directly to AMP, but its effectiveness depends on a finite substrate^9^ and comes at the expense of toxic nucleotide production that interferes with genomic stability once quality control enzymes peak/saturate^22^. Preservation of basal AdN abundance by nucleotide salvage may therefore conceal a vulnerability that becomes apparent when contraction, regeneration or inflammation increases nucleotide turnover; one with the potential to amplify muscle pathology and degradation. ASA functionally tested this constraint. By entering the pathway downstream of ADSS1, ASA increased ATP, ADP and adenine; neither ribose nor DMF reproduced this effect. Additional pentose supply therefore cannot overcome restricted IMP>adenylosuccinate conversion, however it can stimulate nucleotide salvage through shuttling of nucleosides derived from *de novo* IMP synthesis and degradation to AdNs^23^. Fumarate-responsive signalling operates downstream of the nucleotide bottleneck and appears necessary for adaptive mitochondrial coupling to the cytosolic metabolic program. These comparisons establish the control exerted by the ADSS1 reaction even when steady-state ATP remains compensated. They also show that this node has separable outputs: adenylate production depends on bypassing ADSS1, whereas several inflammatory, adipogenic and chromatin-associated programs can be modified without expanding the AdN pool.

### Purine retention and fumarate offset inflammatory stress

Purine escape provides a direct connection between ADSS1 suppression and the inflammatory environment of dystrophic muscle. IMP not retained through the PNC can be degraded through inosine and hypoxanthine to xanthine and urate, a known proinflammatory compound^24^. HPRT-mediated recovery of hypoxanthine conserves purines and limits their delivery to xanthine oxidoreductase (XOR, encoded by *Xdh*), which can generate superoxide and hydrogen peroxide during terminal purine catabolism. Increased HPRT in *mdx* muscle is consistent with pressure to retain this purine fraction.

The response to ribose illustrates what may occur when upstream supply exceeds the capacity of the ADSS1 node. RIB increased *Xdh* expression without expanding the AdN pool. This is consistent with greater pressure on terminal purine disposal when increased ribose-phosphate availability feeds a restricted IMP>AMP pathway. A similar imbalance occurs in severely ADSS-deficient lymphoma cells, where induction of *de novo* synthesis produces IMP accumulation and inosine release rather than balanced adenylate restoration^19^. This connection is particularly relevant to DMD, where XOR activity is increased and its pharmacological inhibition limits contraction-induced force loss^25^. Purine escape from muscle could therefore reinforce an established oxidant and inflammatory circuit. Muscle injury and immune activation would, in turn, increase ATP turnover and purine release, establishing a feed-forward relationship between nucleotide demand, terminal purine catabolism and inflammatory stress.

Reduced passage through the coupled ADSS1–ADSL reactions may also weaken the opposing defence. Fumarate was lower in *mdx* muscle, and fumarate derivatives activate NRF2-dependent cytoprotection and restrain inflammasome signalling^26^, including through succination of gasdermin D^27^. In the same exercise-aggravated *mdx* cohort, DMF suppressed NF-κB signalling, inflammatory chemokines and macrophage infiltration^16^ without expanding the AdN pool here. DMF can therefore replace selected downstream antioxidant and immunoregulatory actions of fumarate but cannot correct the upstream purine constraint. ASA acts closer to the node by supporting ADSL-dependent production of both AMP and fumarate. ADSS1 suppression may consequently influence inflammatory tone from two directions: (1) by increasing the potential for purine escape; and (2) by limiting fumarate-dependent defence.

### ADSS1 connects purine metabolism with adipogenic remodelling

The lipid phenotype extends the influence of the ADSS1 node beyond merely supporting nucleotide abundance. *Mdx* muscle accumulated triacylglycerols and medium chain acylcarnitines and activated an adipogenic transcriptional program before overt adipocyte replacement was detected. A comparable signature is observed in *mdx* plasma^28^. Intramuscular adiposis is a major pathological feature of both progressive DMD^12^ and ADSS1 myopathy^29^, and the 86% directional overlap in the adipogenesis-associated module places lipid remodelling alongside purine maintenance as a shared feature of both disorders.

ASA, ribose and DMF suppressed *Cidea* and *Fabp4* despite producing distinct metabolomic responses and despite only ASA expanding the AdN pool. Adipogenic transcription is therefore determined by downstream mitochondrial carbon handling, fumarate-sensitive signalling or metabolite-dependent chromatin regulation rather nucleotide pools. Longer-term studies support the biological relevance of this early transcriptional effect. ASA reduced muscle adiposis in DMD patients^30^ and older *mdx* mice^31^, while DMF reduced adipogenic remodelling in *mdx* muscle. The present findings extend this response to ribose.

ADSS1 may also connect metabolism with chromatin more directly. In adipocytes, ADSS1 deficiency alters HDAC3 localisation and increases H3K27 acetylation at the glycerol kinase locus, promoting glycerol-dependent fatty-acid re-esterification^11^. This response retains fatty acids in triacylglycerol and supports lipid synthesis–oxidation cycling; its physiological outcome consequently depends on the oxidative capacity and cellular identity of the tissue. In adipose tissue it supports thermogenic cycling^11^, whereas in dystrophic muscle with impaired oxidative handling, a comparable shift could favour lipid retention and contribute to lipidosis. The increase in *Hdac1* in *mdx* muscle, its reduction by ASA and the broader chromatin response to all three interventions are consistent with metabolic–epigenetic coupling, although they do not establish the adipocyte HDAC3 mechanism in muscle *per se*.

### The ADSS node may have systemic metabolic reach

Although the present study is centred on skeletal muscle, the metabolic reach of ADSS regulation may be broader. ADSS2 is widely expressed^32^, while ADSS1 is enriched in skeletal and cardiac muscle but is also expressed at lower levels in other metabolically active tissues^8^. Tissue-selective ADSS1 functions are already evident in the literature. Adipose-specific *Adss1* deletion alters lipid cycling and whole-body energy expenditure, while pancreatic β-cell studies link the ADSS product, adenylosuccinate, with glucose-stimulated insulin secretion^33^. The emerging evidence that ADSS-generated adenylosuccinate also regulates islet size and proliferation^34^ further links this pathway to pancreatic adaptation and systemic glucose control. These observations raise the possibility that coordinated suppression of ADSS1 and ADSS2 could have consequences beyond the affected muscle. If the purine-retention signature identified here also occurs in pancreatic islets, adipose tissue, liver or heart, it could influence insulin secretion, substrate partitioning and whole-body glucose handling. This possibility is pertinent given emerging evidence of systemic glucose dysregulation in *mdx* mice^35,36^. While the present data do not establish a multi-organ ADSS program, they do provide a testable framework that ADSS regulation may couple tissue-specific purine demand to endocrine and systemic metabolic adaptation.

## Limitations of the study

This study provides an integrated view of the molecular architecture surrounding ADSS1 suppression in dystrophic skeletal muscle and resolves distinct responses to interventions positioned around the ADSS1 reaction. However, metabolite abundance and gene expression represent pathway state rather than flux. Stable-isotope tracing will be needed to determine how ADSS1 suppression alters purine synthesis, salvage and escape, and how these changes intersect with TCA-cycle and XOR activity. Likewise, follow up studies require targeted AdN measures for absolute concentrations, calculations of ratios, pool size and energy charge which could not be determined through mass spectrometry approaches. The analyses were also performed in bulk quadriceps, which contains myofibres together with immune, stromal and vascular populations. Single-nucleus and spatial approaches could resolve the cellular origins of the inflammatory, adipogenic and purine-maintenance programs identified here. Furthermore, the cross-species comparison establishes molecular convergence between *mdx* and human ADSS1-myopathy muscle but does not establish ADSS1 suppression as the sole driver of dystrophic metabolism. Likewise, ASA, ribose and DMF were used as metabolically positioned interventions rather than target-selective probes, and their effects may extend beyond the ADSS1 pathway. Finally, the findings derive from one muscle, disease stage and exercise-aggravated context. Studies across disease progression, muscle groups and physiological demand will be required to determine when the compensated purine state becomes limiting and whether the same ADSS-centred program operates in other tissues.

## Conclusions

ADSS1 suppression in dystrophic muscle does not produce immediate resting ATP failure. Rather, muscle preserves AdNs by increasing salvage while both ADSS isoforms decline, free adenine is consumed and the potential for purine escape increases. The shared purine-maintenance program in *mdx* and human ADSS1-myopathy muscle, together with the graded consequences of ADSS deficiency across worms, yeast, lymphoma cells and mice, positions ADSS1 as a deliberately regulated metabolic control node rather than a passive marker of dystropathology. The interventions exposed the outputs controlled by this node and highlight that production of toxic/disruptive nucleotides are a consequence of nucleotide salvage compensation. ASA bypasses the restricted reaction and expands the AdN pool, while fumarate- and pentose-directed interventions modify inflammatory, adipogenic and chromatin-associated programs without necessarily restoring AdN abundance. ADSS1 therefore connects purine retention with fumarate-dependent defence and muscle cell-state remodelling. Its suppression creates a compensated but potentially demand-limited metabolic state – one that is pharmacologically accessible and may have consequences extending beyond skeletal muscle.

## Resource availability

### Lead contact

Requests for further information and resources should be directed to and will be fulfilled by the lead contact, Emma Rybalka.

### Materials availability

This study did not generate new unique reagents.

### Data and code availability

Raw and processed data files are available in public repositories.

## Acknowledgments

Funding sources comprise philanthropic donations from The Estate of Charles A. Bonsett (Dystrophy Concepts, E.R.), and grants from the USA Muscular Dystrophy Association (MDA871929, E.R.) and Australian Physiological Society (S.K.). The authors wish to thank Professor Alan Beggs and Dr Behzad Moghadaszadeh for donation of ADSS1 and ADSS2 antibodies; Pat Quiambao, for laboratory assistance at Metabolomics Australia, A/Prof Jujiao Kuang for laboratory assistance at Victoria University, and Ms Tricia Murphy and Dr Steven Holloway for support at Victoria University Animal Facilities.

## Author contributions

Conceptualization, E.R., D.F. and C.A.T; methodology, E.R., S.K., D.G.C., B.N., D.D-L., D.F. and C.A.T.; investigation, E.R., S.K., B.Q., D.G.C., A.T., N.K., A.P., B.N., D.D-L., D.F., H.J.P., and C.A.T.; funding acquisition, E.R. and S.K.; project management; E.R., S.K. and C.A.T.; writing – original draft, E.R. and S.K.; writing- review and editing, E.R., S.K., B.Q., D.G.C., A.T., G.S.M., N.K., A.P., C.G.S., B.N., D.D-L., D.F., J.J.B., A.L. H.J.P. and C.A.T.

## Methods

### Study Approval and Animal Treatment

This study was approved by the Victoria University Animal Ethics Committee (20/005 and 20/006) and complied with the Australian Code of Practice for Care and Use of Animals for Scientific Purposes. Male C57BL/10ScSnJ (WT) and dystrophin-deficient C57BL/10ScSn- *Dmd^mdx^/*J (*mdx*) mice were bred at Western Centre for Health, Research and Education (Sunshine Hospital, Victoria, Australia) and housed under a 12-hour light-dark cycle in 20- 25°C and 40% humidity conditions. At 21 days of age, male mice were weaned and randomly allocated to treatment groups. Using 0.5% methylcellulose vehicle (VEH), *mdx* mice were treated daily by oral gavage at a dose rate of 325mg/kg/day ASA, 100mg/kg/day DMF or 1.6g/kg/day ribose. WT and *mdx* control groups received daily VEH gavage (10% v/b.w.). Food and water consumption and anthropometry were normal across the 5-week treatment period^16^. From 4 weeks of age, mice were exercised bi-weekly via treadmill (30 min, 12 m/min), which exacerbated murine DMD as previously published^16^.

### Surgical Procedure

At the experimental endpoint of 8 weeks of age, precisely 2 days after the final treadmill run, mice were weighed, dosed with their final treatment, and then deeply anaesthetised with isoflurane (4% induction and 2.5% maintenance). The left quadriceps were excised and immediately snap frozen for transcriptomic, metabolomic and lipidomic analyses. The right quadriceps were snap frozen and then later slow thawed at 4°C and fixed in 10% neutral buffered formalin for 48 hours. Samples were subsequently transferred to 70% ethanol until paraffin embedding and were then processed for immunohistochemical analysis.

### Western Blot

Primary antibodies: anti-ADSS1 and ADSS2 (1:1000; Boston Children’s Hospital, Beggs Lab), anti-ADSL (1:1000; PA5-87207; ThermoFisher), anti-AMPD1 (1;1000; NBP2-24509; Novus Biologicals) diluted in bovine serum albumin were used. All membranes were incubated with a horseradish peroxidase-conjugated secondary antibody (anti-rabbit IgG; Vector Laboratories; 1:5000) diluted in 3% non-fat milk for 1 hour at room temperature, followed by imaging, Coomassie blue staining and normalisation to total protein.

### Transcriptomics

RNA sequencing and transcriptomic analysis was performed as previously detailed16.

### Metabolomics and Lipidomics

Metabolomics and lipidomics analyses were conducted at Metabolomics Australia core (Bio21 Molecular Science and Biotechnology Institute, University of Melbourne, Parkville, Australia). Thirty mg of snap-frozen quadriceps (*n* = 8 per group) were homogenised using a handheld homogeniser (6 x 10 seconds blocks at 13,000 rpm) in 300 μl of ice-cold 3:1 Methanol:Milli-Q water containing MA-Internal Standards. Extracts were vortexed for 10 seconds, mixed on a thermomixer and then centrifuged at 4°C for 10 minutes at maximum speed to produce the supernatant. Polar metabolite analysis was performed on the Orbitrap ID-X Tribrid mass spectrometer (Thermo Scientific, Waltham, MA, USA) coupled to a Vanquish Horizon UHPLC system (Thermo Scientific, Waltham, MA, USA) using 100 μl of sample supernatant. The pooled biological quality control sample (pbQc) was generated using 20 μl of each sample and run after every five-biological samples.

#### Targeted lipid analysis

The remaining homogenate and lysate were further extracted in 300 μl of 100% chloroform (containing MA-lipid Internal Standards) was added to create a ratio of 2:1 chloroform:methanol for the remaining homogenate. Extracts were vigorously vortexed to resuspend the tissue pellet. Samples were mixed on a thermomixer and centrifuged at 10°C for 5 mins at 25,155 x g. The LCMS methodologies for polar metabolite and targeted lipid analyses are outlined previously^37^.

### Bioinformatics

Statistical data analysis and data normalisation of metabolomics and lipidomics was performed in MetaboAnalyst 6.0. Data were normalised by median log transformation without data scaling. For metabolomics and lipidomics, a 1.0 log_2_ fold change threshold and a raw *p* value of <0.05 was applied to distinguish significantly changed metabolites/lipid species.

For transcriptomics, differential expression analyses were conducted using Degust and a 1.0 log_2_ fold change threshold and raw *p* value of <0.05 was considered significant. Pathways enrichment was performed using Reactome as previously described^16^.

For metabolomic, lipidomic and transcriptomic heatmaps, unbiased log_2_ fold change is reported and a raw *p* value of <0.05 was considered significant. For metabolite pathways analysis, a raw *p* value of <0.05 was considered significant with integrative pathways analysis significance criteria set as at least one metabolite and gene hit and an impact score >0.2.

## References

1. Park, H.J., Hong, Y.B., Choi, Y.C., Lee, J., Kim, E.J., Lee, J.S., Mo, W.M., Ki, S.M., Kim, H.I., Kim, H.J., et al. (2016). ADSSL1 mutation relevant to autosomal recessive adolescent onset distal myopathy. Ann Neurol 79, 231–243. 10.1002/ana.24550.

2. Bakay, M., Zhao, P., Chen, J., and Hoffman, E.P. (2002). A web-accessible complete transcriptome of normal human and DMD muscle. Neuromuscul Disord 12 *Suppl 1*, S125–141. 10.1016/s0960-8966(02)00093-7.

3. Capitanio, D., Moriggi, M., Torretta, E., Barbacini, P., De Palma, S., Vigano, A., Lochmuller, H., Muntoni, F., Ferlini, A., Mora, M., and Gelfi, C. (2020). Comparative proteomic analyses of Duchenne muscular dystrophy and Becker muscular dystrophy muscles: changes contributing to preserve muscle function in Becker muscular dystrophy patients. J Cachexia Sarcopenia Muscle 11, 547–563. 10.1002/jcsm.12527.

4. Tsitsipatis, D., Mazan-Mamczarz, K., Si, Y., Herman, A.B., Yang, J.H., Guha, A., Piao, Y., Fan, J., Martindale, J.L., Munk, R., et al. (2022). Transcriptomic analysis of human ALS skeletal muscle reveals a disease-specific pattern of dysregulated circRNAs. Aging (Albany NY) 14, 9832–9859. 10.18632/aging.204450.

5. Xu, X., Yang, Q., Liu, Z., Zhang, R., Yu, H., Wang, M., Chen, S., Xu, G., Shao, Y., and Le, W. (2023). Integrative analysis of metabolomics and proteomics unravels purine metabolism dysregulation in the SOD1(G93A) mouse model of amyotrophic lateral sclerosis. Neurobiol Dis 181, 106110. 10.1016/j.nbd.2023.106110.

6. Strand, A.D., Aragaki, A.K., Shaw, D., Bird, T., Holton, J., Turner, C., Tapscott, S.J., Tabrizi, S.J., Schapira, A.H., Kooperberg, C., and Olson, J.M. (2005). Gene expression in Huntington’s disease skeletal muscle: a potential biomarker. Human Molecular Genetics 14, 1863–1876. 10.1093/hmg/ddi192.

7. Lowenstein, J.M., and Goodman, M.N. (1978). The purine nucleotide cycle in skeletal muscle. Fed Proc 37, 2308–2312.

8. Li, X., Zhu, Z., Mo, D., Wang, H., Yang, S., Zhao, S., and Li, K. (2007). Comparative molecular characterization of ADSS1 and ADSS2 genes in pig (Sus scrofa). Comp Biochem Physiol B Biochem Mol Biol 147, 271–277. 10.1016/j.cbpb.2007.01.013.

9. Van den Berghe, G., Bontemps, F., Vincent, M.F., and Van den Bergh, F. (1992). The purine nucleotide cycle and its molecular defects. Prog Neurobiol 39, 547–561. 10.1016/0301-0082(92)90006-z.

10. Van den Berghe, G., Vincent, M.F., and Jaeken, J. (1997). Inborn errors of the purine nucleotide cycle: adenylosuccinase deficiency. J Inherit Metab Dis 20, 193–202. 10.1023/a:1005304722259.

11. Sun, J., Alimujiang, M., Li, W., Chen, S., Su, Y., Hu, T., Lu, X., Ye, Y., Bai, N., Hu, F., et al. (2026). Adenylosuccinate Synthase 1 Deficiency Improves Energy Metabolism by Promoting Adipose Tissue Re-esterification via Glycerol Kinase Upregulation. Adv Sci (Weinh) 13, e06270. 10.1002/advs.202506270.

12. Duan, D., Goemans, N., Takeda, S., Mercuri, E., and Aartsma-Rus, A. (2021). Duchenne muscular dystrophy. Nat Rev Dis Primers 7, 13. 10.1038/s41572-021-00248-3.

13. Timpani, C.A., Hayes, A., and Rybalka, E. (2015). Revisiting the dystrophin-ATP connection: How half a century of research still implicates mitochondrial dysfunction in Duchenne Muscular Dystrophy aetiology. Med Hypotheses 85, 1021–1033. 10.1016/j.mehy.2015.08.015.

14. Onopiuk, M., Brutkowski, W., Wierzbicka, K., Wojciechowska, S., Szczepanowska, J., Fronk, J., Lochmüller, H., Górecki, D.C., and Zabłocki, K. (2009). Mutation in dystrophin-encoding gene affects energy metabolism in mouse myoblasts. Biochem Biophys Res Commun 386, 463–466. 10.1016/j.bbrc.2009.06.053.

15. Park, H.J., Hong, J.M., Lee, J.H., Shin, H.Y., Kim, S.M., Park, K.D., Lee, J.H., and Choi, Y.C. (2019). Comparative transcriptome analysis of skeletal muscle in ADSSL1 myopathy. Neuromuscul Disord 29, 274–281. 10.1016/j.nmd.2018.11.003.

16. Kourakis, S., Timpani, C.A., Bagaric, R.M., Qi, B., Ali, B.A., Boyer, R., Spiesberger, G., Kandhari, N., Yan, X., Kuang, J., et al. (2025). Repurposed Nrf2 activator dimethyl fumarate rescues muscle inflammation and fibrosis in an aggravated mdx mouse model of Duchenne muscular dystrophy. Redox Biol 84, 103676. 10.1016/j.redox.2025.103676.

17. Liu, G., Lee, D.P., Schmidt, E., and Prasad, G.L. (2019). Pathway Analysis of Global Metabolomic Profiles Identified Enrichment of Caffeine, Energy, and Arginine Metabolism in Smokers but Not Moist Snuff Consumers. Bioinform Biol Insights 13, 1177932219882961. 10.1177/1177932219882961.

18. Patil, R.R., Franklin, L.P., Jin, M.D., and Hanna-Rose, W. (2025). Adenylosuccinate synthetase deficiency and purine nucleotide cycle disruption impair neuromuscular function in Caenorhabditis elegans. Molecular Genetics and Metabolism 146, 109261. 10.1016/j.ymgme.2025.109261.

19. Ullman, B., Wormsted, M.A., Cohen, M.B., and Martin, D.W., Jr. (1982). Purine oversecretion in cultured murine lymphoma cells deficient in adenylosuccinate synthetase: genetic model for inherited hyperuricemia and gout. Proc Natl Acad Sci U S A 79, 5127–5131. 10.1073/pnas.79.17.5127.

20. Tarakhovskaya, E.R., Andreychuk, Y.V., Bilova, T.E., Wiesner, C., Pavlov, Y.I., and Stepchenkova, E.I. (2025). Vulnerable Nucleotide Pools and Genomic Instability in Yeast Strains with Deletion of the ADE12 Gene Encoding for Adenylosuccinate Synthetase. Int J Mol Sci 26. 10.3390/ijms26083458.

21. Kim, M.E., Yammine, K.M., Hickey, E.T., Matias, C., Dubosclard, L.C., Widrick, J.J., Brault, J.J., Moghadaszadeh, B., and Beggs, A.H. (2025). Loss of adenylosuccinate synthetase 1 in mice recapitulates features of ADSS1 myopathy. Hum Mol Genet. 10.1093/hmg/ddaf167.

22. Wang, P., Wang, C., and Wang, Y. (2026). Nucleotide salvage, genome instability, and potential therapeutic applications. Nucleic Acids Res 54. 10.1093/nar/gkag099.

23. Brault, J.J., and Terjung, R.L. (2001). Purine salvage to adenine nucleotides in different skeletal muscle fiber types. J Appl Physiol (1985) 91, 231–238. 10.1152/jappl.2001.91.1.231.

24. Bortolotti, M., Polito, L., Battelli, M.G., and Bolognesi, A. (2021). Xanthine oxidoreductase: One enzyme for multiple physiological tasks. Redox Biology 41, 101882. 10.1016/j.redox.2021.101882.

25. Lindsay, A., McCourt, P.M., Karachunski, P., Lowe, D.A., and Ervasti, J.M. (2018). Xanthine oxidase is hyper-active in Duchenne muscular dystrophy. Free Radic Biol Med 129, 364–371. 10.1016/j.freeradbiomed.2018.10.404.

26. Kourakis, S., Timpani, C.A., de Haan, J.B., Gueven, N., Fischer, D., and Rybalka, E. (2021). Targeting Nrf2 for the treatment of Duchenne Muscular Dystrophy. Redox Biol 38, 101803. 10.1016/j.redox.2020.101803.

27. Humphries, F., Shmuel-Galia, L., Ketelut-Carneiro, N., Li, S., Wang, B., Nemmara, V.V., Wilson, R., Jiang, Z., Khalighinejad, F., Muneeruddin, K., et al. (2020). Succination inactivates gasdermin D and blocks pyroptosis. Science 369, 1633–1637. 10.1126/science.abb9818.

28. Tsonaka, R., Seyer, A., Aartsma-Rus, A., and Spitali, P. (2021). Plasma lipidomic analysis shows a disease progression signature in mdx mice. Sci Rep 11, 12993. 10.1038/s41598-021-92406-6.

29. Motoda, A., Takahashi, T., Watanabe, C., Tachiyama, Y., Ochi, K., Saito, Y., Iida, A., Nishino, I., and Maruyama, H. (2021). An autopsied case of ADSSL1 myopathy. Neuromuscul Disord 31, 1220–1225. 10.1016/j.nmd.2021.07.011.

30. Bonsett, C.A., and Rudman, A. (1992). The dystrophin connection--ATP? Med Hypotheses 38, 139–154. 10.1016/0306-9877(92)90087-s.

31. Timpani, C.A., Goodman, C.A., Stathis, C.G., White, J.D., Mamchaoui, K., Butler-Browne, G., Gueven, N., Hayes, A., and Rybalka, E. (2020). Adenylosuccinic acid therapy ameliorates murine Duchenne Muscular Dystrophy. Sci Rep 10, 1125. 10.1038/s41598-020-57610-w.

32. Tran, D.H., Kim, D., Kesavan, R., Brown, H., Dey, T., Soflaee, M.H., Vu, H.S., Tasdogan, A., Guo, J., Bezwada, D., et al. (2024). *De novo* and salvage purine synthesis pathways across tissues and tumors. Cell 187, 3602–3618.e3620. 10.1016/j.cell.2024.05.011.

33. Gooding, J.R., Jensen, M.V., Dai, X., Wenner, B.R., Lu, D., Arumugam, R., Ferdaoussi, M., MacDonald, P.E., and Newgard, C.B. (2015). Adenylosuccinate Is an Insulin Secretagogue Derived from Glucose-Induced Purine Metabolism. Cell Rep 13, 157–167. 10.1016/j.celrep.2015.08.072.

34. Inoue, R., Tsuno, T., Nishimura, T., Fukushima, S., Hirai, S., Shimoda, M., Yoshinari, Y., Sakai, C., Kin, T., Lim, E.X.H., et al. (2025). Adenylosuccinate Mediates Imeglimin-Induced Proliferative and Antiapoptotic Effects in β-Cells. Diabetes 74, 1589–1602. 10.2337/db24-1090.

35. Major, G.S., Timpani, C.A., Lalunio, H., Chen, J., Boatner, L., Giourmas, N., Kourakis, S., van den Berg, E., Salimova, E., Eliades, J., et al. (2026). Stress unmasks impaired endocrine coordination of glucose metabolism in dystrophin deficiency. bioRxiv, 2026.2002.2022.707291. 10.64898/2026.02.22.707291.

36. Podkalicka, P., Mucha, O., Kaziród, K., Szade, K., Stępniewski, J., Ivanishchuk, L., Hirao, H., Pośpiech, E., Józkowicz, A., Kupiec-Weglinski, J.W., et al. (2022). miR- 378 affects metabolic disturbances in the mdx model of Duchenne muscular dystrophy. Sci Rep 12, 3945. 10.1038/s41598-022-07868-z.

37. Lu, T., Freytag, L., Narayana, V.K., Moore, Z., Oliver, S.J., Valkovic, A., Nijagal, B., Peterson, A.L., de Souza, D.P., McConville, M.J., et al. (2023). Matrix Selection for the Visualization of Small Molecules and Lipids in Brain Tumors Using Untargeted MALDI-TOF Mass Spectrometry Imaging. Metabolites 13. 10.3390/metabo13111139.

