## Supplemental Data for "ADSS1 is a suppressed metabolic control node in dystrophic skeletal muscle"

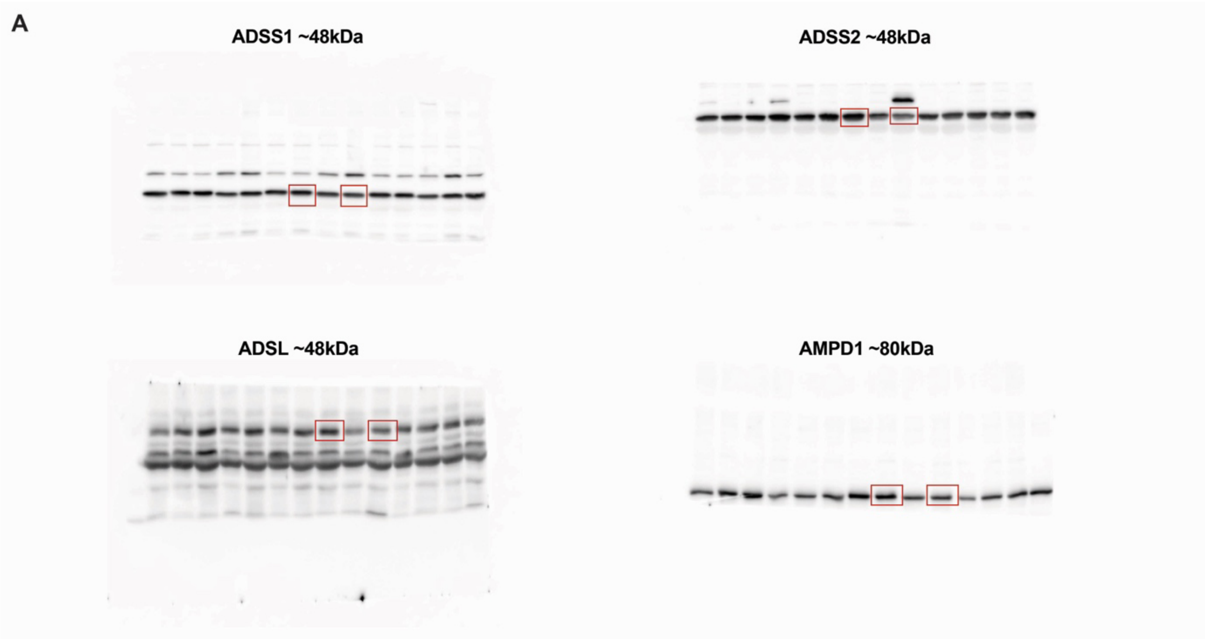

**Figure S1. Full-length blots of muscle ADSS1, ADSS2, ADSL and AMPD1 protein expression supplemental to Figure 1.**

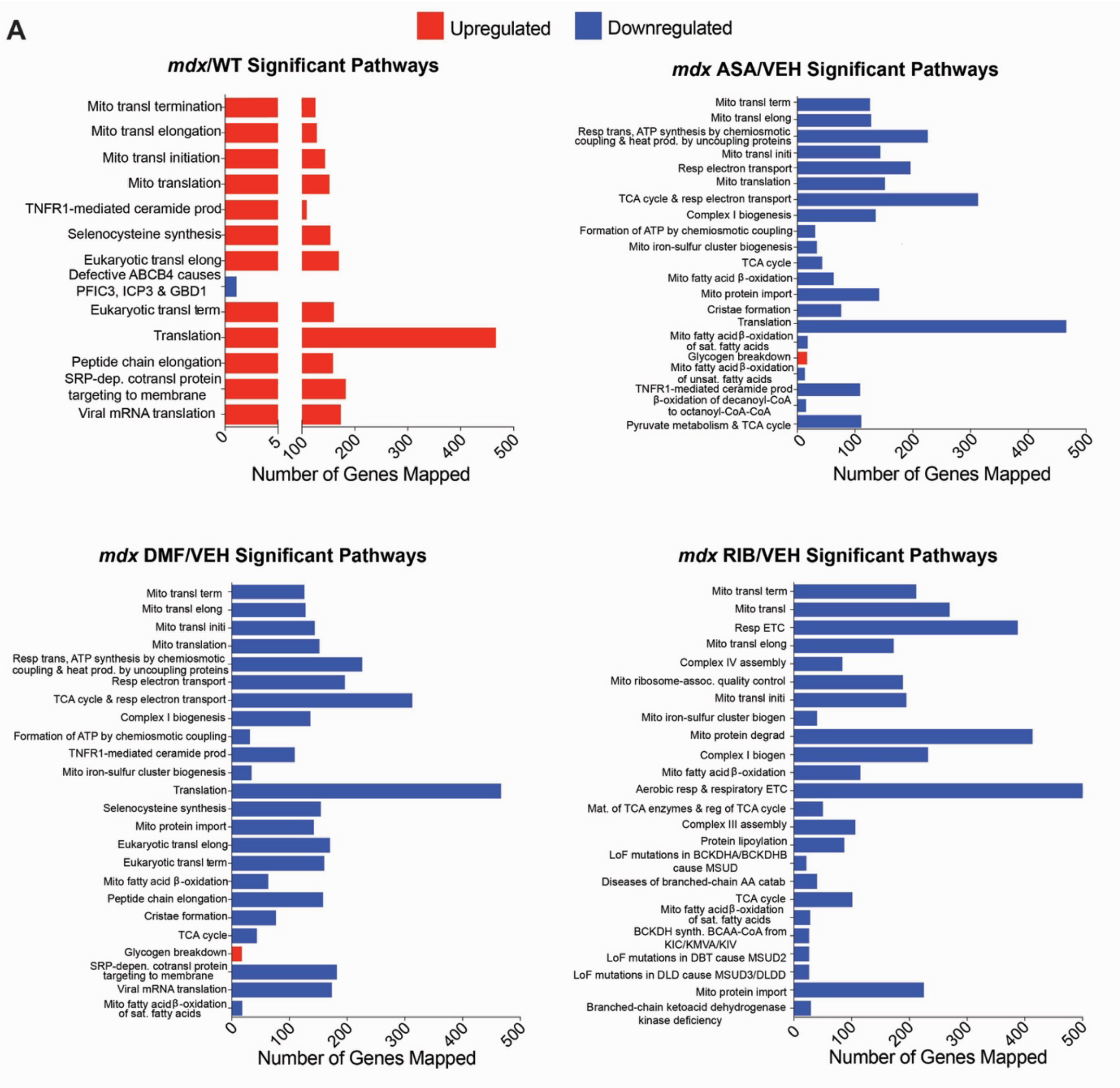

**Figure S2. Significantly regulated muscle transcriptomic pathways displayed as number of genes mapped.**

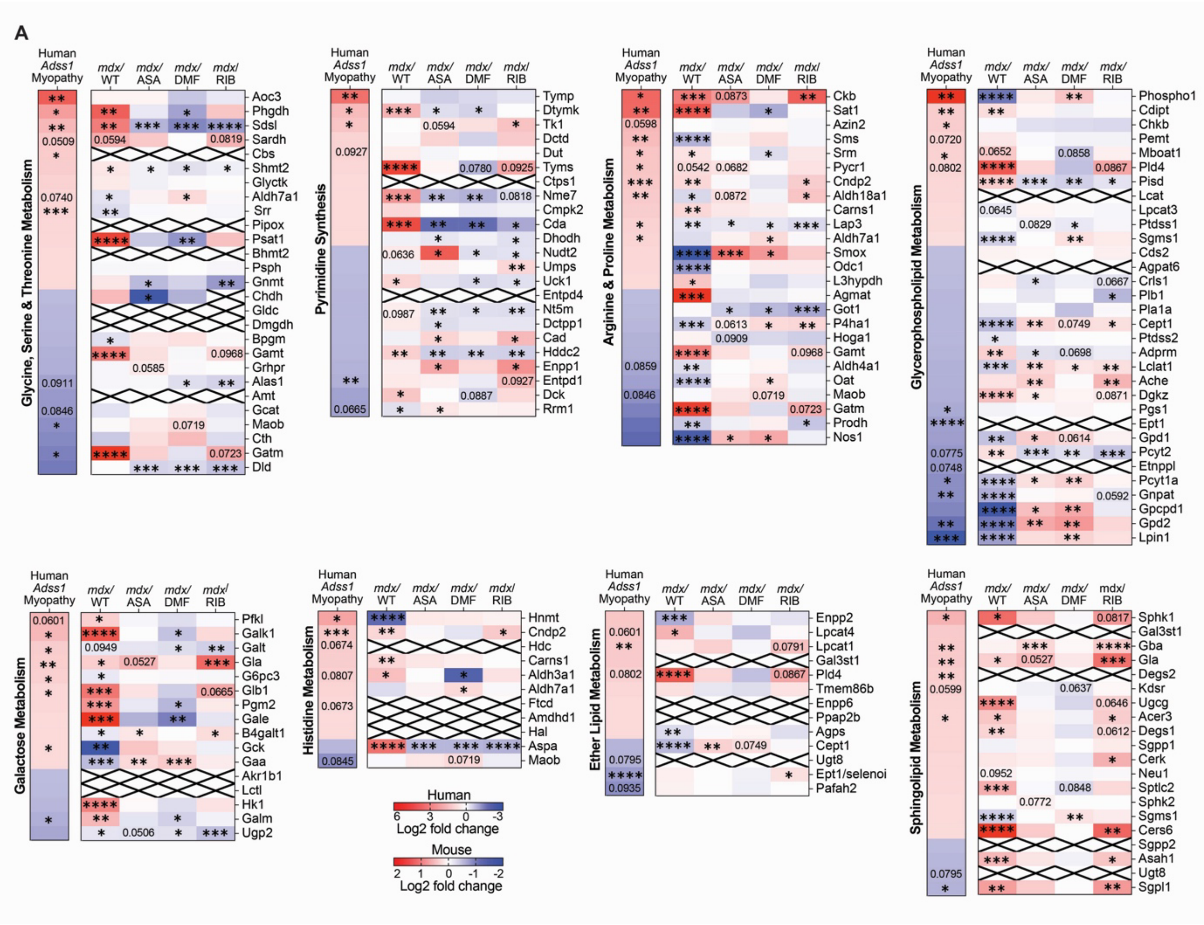

**Figure S3. Cross-species transcriptomic overlay of the most impacted integrated metabolic pathways in *mdx* muscle compared with human primary ADSS1 deficiency and modulatory capacity of adenylosuccinic acid (ASA), dimethyl fumarate (DMF) and ribose (RIB).** *$p$<0.05, **$p$<0.01, ***$p$<0.001, ****$p$<0.0001 different to control. Heatmap crosses denote the gene was not detected in a dataset. *P* values are reported for trends ($p$=>0.05, <0.01).

#### Supplementary Table 1. Significantly upregulated and downregulated metabolites

| *mdx*/WT | | | *mdx* ASA/VEH | | | *mdx* DMF/VEH | | | *mdx* RIB/VEH | | |
|---|---|---|---|---|---|---|---|---|---|---|---|
| Metabolite | log2(FC) | -log$_{10}$($p$) | Metabolite | log2(FC) | -log$_{10}$($p$) | Metabolite | log2(FC) | -log$_{10}$($p$) | Metabolite | log2(FC) | -log$_{10}$($p$) |
| Kynurenine | 4.386 | 12.71 | Erythrose 4-phosphate | 6.073 | 3.00 | Hypoxanthine | -1.172 | 1.644 | Erythrose 4-phosphate | 4.465 | 1.671 |
| Anthranilate/2-Aminobenzoic acid | 3.311 | 9.624 | Taurocholic acid | 2.839 | 2.096 | Fructose 6-phosphate | -1.474 | 1.754 | Taurocholic acid | 2.711 | 2.419 |
| Inosine triphosphate | 2.869 | 1.538 | Coenzyme A | 2.399 | 5.603 | Glucose 6-phosphate | -1.853 | 2.012 | Xylitol | 2.179 | 7.998 |
| O-Phosphoethanolamine | 1.558 | 5.588 | Acetyl-CoA | 1.573 | 4.709 | Erythrose 4-phosphate | -4.718 | 1.776 | N-Acetylglutamine | 1.906 | 2.317 |
| Glycerophosphocholine | 1.333 | 4.934 | Glucosamine | 1.499 | 3.812 | | | | Taurodeoxycholic acid | 1.627 | 1.481 |
| Stachyose | 1.243 | 2.030 | N-Acetylglutamine | 1.344 | 2.561 | | | | Acetylglycine | 1.560 | 2.051 |
| Deoxycholic acid | 1.221 | 1.581 | Glycerol | 1.343 | 2.971 | | | | N-Acetylglutamic acid | 1. | 3. |

| | | | | | | | | | |
|---|---|---|---|---|---|---|---|---|---|
| | | | | | | | | 493 | 104 |
| beta-Glycerophosphoric acid | 1.166 | 3.982 | NADH | 1.306 | 3.591 | | N-Acetyltyrosine | 1.281 | 1.368 |
| Maltotriose | 1.038 | 1.880 | Adenosine triphosphate | 1.290 | 3.018 | | Succinic acid | 1.203 | 2.222 |
| Raffinose | 1.037 | 1.661 | Succinic acid | 1.271 | 2.423 | | Coenzyme A | 1.109 | 1.807 |
| Orotidine | 1.036 | 5.668 | Glycerol 3-phosphate | 1.188 | 1.695 | | Orotic acid | 1.041 | 1.950 |
| NADH | -1.043 | 2.617 | Inosine monophosphate | 1.121 | 3.121 | | Cortexolone | 1.033 | 1.855 |
| Proline | -1.076 | 4.264 | N-Acetylglutamic acid | 1.036 | 2.769 | | Levulinic acid | -1.008 | 3.493 |
| Glycine | -1.101 | 4.393 | Guanosine monophosphate | -1.092 | 1.415 | | Stachyose | -1.414 | 2.250 |
| Carnosine | -1.177 | 6.368 | Ribulose 5-phosphate | -1.347 | 2.824 | | Maltose | -1.421 | 2.181 |

| | | | | | | | | | |
|---|---|---|---|---|---|---|---|---|---|
| Asparagine | -1.208 | 2.663 | 2-Ketobutyric acid | -1.506 | 3.724 | | Maltotriose | -1.597 | 2.889 |
| 3-Phenylbutyric acid | -1.241 | 1.579 | Maltose | -1.645 | 2.661 | | Raffinose | -1.603 | 2.388 |
| Vanillin | -1.333 | 1.723 | Stachyose | -1.745 | 3.168 | | | | |
| Acetylglycine | -1.395 | 4.212 | Adenylosuccinic acid | -1.823 | 4.172 | | | | |
| Isoeugenol | -1.434 | 3.379 | Maltotriose | -1.957 | 3.996 | | | | |
| Adenine | -1.604 | 1.798 | Uridine 5'-monophosphate | -2.287 | 2.671 | | | | |
| 4-Hydroxyproline | -1.647 | 4.432 | Raffinose | -2.478 | 4.308 | | | | |
| 5-Phenylvaleric acid | -1.718 | 3.250 | 3-Phenylbutyric acid | -3.279 | 1.369 | | | | |

#### Supplementary Table 2. Metabolites Pathway Analysis *mdx*/WT

| Pathway | Total metabolites in pathway | Hits | Raw *p* value | -log10(*p*) | Impact |
|---|---|---|---|---|---|
| **Significant Pathways** | | | | | |
| Ammonia Recycling | 25 | 3 | 0.004 | 2.383 | 0.177 |
| Galactose Metabolism | 31 | 3 | 0.008 | 2.114 | 0.009 |
| Purine Metabolism | 63 | 4 | 0.009 | 2.059 | 0.050 |
| Ethanol Degradation | 15 | 2 | 0.017 | 1.768 | 0.515 |
| Carnitine Synthesis | 16 | 2 | 0.019 | 1.713 | 0.000 |
| Tryptophan Metabolism | 55 | 3 | 0.037 | 1.435 | 0.098 |
| Bile Acid Biosynthesis | 60 | 3 | 0.046 | 1.338 | 0.032 |
| Beta-Alanine Metabolism | 26 | 2 | 0.048 | 1.315 | 0.190 |
| **Non-Significant Pathways** | | | | | |
| Retinol Metabolism | 30 | 2 | 0.063 | 1.202 | 0.082 |
| Pyruvate Metabolism | 30 | 2 | 0.063 | 1.202 | 0.415 |
| Aspartate Metabolism | 34 | 2 | 0.078 | 1.106 | 0.250 |
| Histidine Metabolism | 35 | 2 | 0.083 | 1.084 | 0.183 |
| Porphyrin Metabolism | 36 | 2 | 0.087 | 1.062 | 0.000 |
| Malate-Aspartate Shuttle | 7 | 1 | 0.094 | 1.028 | 0.000 |
| Methionine Metabolism | 39 | 2 | 0.100 | 1.002 | 0.009 |
| Glycerol Phosphate Shuttle | 8 | 1 | 0.107 | 0.973 | 0.429 |
| Glucose-Alanine Cycle | 9 | 1 | 0.119 | 0.924 | 0.000 |
| De Novo Triacylglycerol Biosynthesis | 9 | 1 | 0.119 | 0.924 | 0.000 |
| Glutamate Metabolism | 45 | 2 | 0.127 | 0.897 | 0.058 |
| Arginine and Proline Metabolism | 48 | 2 | 0.141 | 0.850 | 0.097 |
| Cardiolipin Biosynthesis | 11 | 1 | 0.144 | 0.843 | 0.000 |
| Glycine and Serine Metabolism | 50 | 2 | 0.151 | 0.821 | 0.115 |
| Ketone Body Metabolism | 12 | 1 | 0.156 | 0.808 | 0.000 |
| Threonine and 2-Oxobutanoate Degradation | 13 | 1 | 0.168 | 0.776 | 0.000 |
| Phosphatidylethanolamine Biosynthesis | 13 | 1 | 0.168 | 0.776 | 0.500 |
| Alanine Metabolism | 14 | 1 | 0.179 | 0.746 | 0.000 |
| Mitochondrial Electron Transport Chain | 15 | 1 | 0.191 | 0.719 | 0.156 |
| Butyrate Metabolism | 16 | 1 | 0.202 | 0.694 | 0.000 |
| Nucleotide Sugars Metabolism | 16 | 1 | 0.202 | 0.694 | 0.000 |
| Mitochondrial Beta-Oxidation of Short Chain Saturated Fatty Acids | 17 | 1 | 0.214 | 0.670 | 0.080 |
| Betaine Metabolism | 18 | 1 | 0.225 | 0.648 | 0.000 |
| Pterine Biosynthesis | 18 | 1 | 0.225 | 0.648 | 0.000 |
| Transfer of Acetyl Groups into Mitochondria | 18 | 1 | 0.225 | 0.648 | 0.043 |
| Phosphatidylcholine Biosynthesis | 18 | 1 | 0.225 | 0.648 | 0.220 |
| Phytanic Acid Peroxisomal Oxidation | 19 | 1 | 0.236 | 0.627 | 0.000 |

| | | | | | |
|---|---|---|---|---|---|
| Glutathione Metabolism | 19 | 1 | 0.236 | 0.627 | 0.061 |
| Glycolysis | 20 | 1 | 0.247 | 0.608 | 0.000 |
| Lysine Degradation | 20 | 1 | 0.247 | 0.608 | 0.093 |
| Estrone Metabolism | 20 | 1 | 0.247 | 0.608 | 0.160 |
| Caffeine Metabolism | 20 | 1 | 0.247 | 0.608 | 0.250 |
| Mitochondrial Beta-Oxidation of Medium Chain Saturated Fatty Acids | 22 | 1 | 0.268 | 0.572 | 0.150 |
| Glycerolipid Metabolism | 23 | 1 | 0.278 | 0.555 | 0.000 |
| Urea Cycle | 23 | 1 | 0.278 | 0.555 | 0.000 |
| Androstenedione Metabolism | 23 | 1 | 0.278 | 0.555 | 0.161 |
| Cysteine Metabolism | 24 | 1 | 0.289 | 0.540 | 0.000 |
| Folate Metabolism | 24 | 1 | 0.289 | 0.540 | 0.005 |
| Mitochondrial Beta-Oxidation of Long Chain Saturated Fatty Acids | 24 | 1 | 0.289 | 0.540 | 0.060 |
| Phospholipid Biosynthesis | 25 | 1 | 0.299 | 0.525 | 0.000 |
| Citric Acid Cycle | 25 | 1 | 0.299 | 0.525 | 0.197 |
| Starch and Sucrose Metabolism | 26 | 1 | 0.309 | 0.510 | 0.000 |
| Fructose and Mannose Degradation | 28 | 1 | 0.328 | 0.484 | 0.000 |
| Androgen and Estrogen Metabolism | 29 | 1 | 0.338 | 0.471 | 0.104 |
| Gluconeogenesis | 30 | 1 | 0.348 | 0.459 | 0.112 |
| Amino Sugar Metabolism | 31 | 1 | 0.357 | 0.447 | 0.000 |
| Nicotinate and Nicotinamide Metabolism | 32 | 1 | 0.366 | 0.436 | 0.079 |
| Fatty Acid Biosynthesis | 33 | 1 | 0.375 | 0.426 | 0.001 |
| Fatty Acid Elongation In Mitochondria | 33 | 1 | 0.375 | 0.426 | 0.411 |
| Sphingolipid Metabolism | 36 | 1 | 0.402 | 0.396 | 0.000 |
| Propanoate Metabolism | 36 | 1 | 0.402 | 0.396 | 0.117 |
| Fatty Acid Metabolism | 40 | 1 | 0.436 | 0.361 | 0.292 |
| Steroidogenesis | 42 | 1 | 0.452 | 0.345 | 0.024 |
| Valine, Leucine and Isoleucine Degradation | 51 | 1 | 0.520 | 0.284 | 0.090 |
| Tyrosine Metabolism | 55 | 1 | 0.548 | 0.262 | 0.097 |

#### Supplementary Table 3. Top 50 Significantly upregulated and downregulated genes.

| *mdx*/WT | | | *mdx* ASA/VEH | | | *mdx* DMF/VEH | | | *mdx* RIB/VEH | | |
|---|---|---|---|---|---|---|---|---|---|---|---|
| Gene name | log2(FC) | $-\log_{10}(p)$ | Gene name | log2(FC) | $-\log_{10}(p)$ | Gene name | log2(FC) | $-\log_{10}(p)$ | Gene name | log2(FC) | $-\log_{10}(p)$ |
| Mdfi | 6.019 | 10.377 | Mmp12 | 3.638 | 1.861 | Igfn1 | 2.896 | 3.122 | Mmp12 | 4.989 | 3.863 |
| Gm48395 | 5.934 | 9.353 | Gpnmb | 3.369 | 3.174 | Gm7265 | 2.861 | 3.909 | Gpnmb | 4.519 | 4.645 |
| Mmp12 | 5.687 | 2.864 | Npy | 3.2008 | 2.177 | A630019I02Rik | 2.635 | 4.015 | Atp6v0d2 | 4.379 | 3.151 |
| Krt8 | 5.647 | 9.604 | Atp6v0d2 | 3.173 | 1.635 | Zbed6 | 2.390 | 4.323 | Gm20056 | 4.269 | 4.771 |
| Ccl7 | 5.530 | 5.086 | Itgax | 3.098 | 1.845 | Prima1 | 2.202 | 3.552 | Itgax | 4.242 | 3.183 |
| Cdkn2a | 5.402 | 9.188 | Hoxd10 | 3.052 | 3.968 | Amd2 | 2.171 | 2.950 | Npy | 4.121 | 3.402 |
| Timp1 | 5.127 | 6.287 | Calm4 | 2.953 | 2.873 | Cdh4 | 2.065 | 2.881 | Tm4sf19 | 3.931 | 2.795 |
| Sln | 5.098 | 7.081 | Tm4sf19 | 2.899 | 2.312 | Odf3l2 | 2.038 | 2.618 | Mcoln3 | 3.744 | 3.496 |
| Tnni1 | 4.996 | 4.837 | Gm13938 | 2.896 | 1.887 | Gm26604 | 1.866 | 4.320 | F7 | 3.678 | 2.311 |
| Prune2 | 4.943 | 11.118 | Mmp9 | 2.884 | 1.560 | 1700001O22Rik | 1.845 | 1.486 | Il7r | 3.656 | 2.443 |
| Ccl8 | 4.941 | 5.550 | Gm20056 | 2.833 | 2.092 | Nt5c1a | 1.824 | 3.212 | Mmp9 | 3.622 | 1.568 |
| Stac2 | 4.930 | 7.704 | Il7r | 2.350 | 1.308 | Fam83d | 1.821 | 2.387 | Tnn | 3.437 | 3.533 |
| Serpina3n | 4.879 | 5.408 | Igfn1 | 2.332 | 3.869 | Doc2b | 1.766 | 2.234 | Creg2 | 3.313 | 2.521 |
| Cryzl2 | 4.798 | 8.647 | Fbn2 | 2.287 | 2.405 | Gm11752 | 1.742 | 1.327 | Spp1 | 3.249 | 2.675 |
| Trem2 | 4.780 | 4.208 | Fat3 | 2.257 | 3.342 | B230312C02Rik | 1.706 | 1.338 | Atp1a3 | 3.237 | 2.534 |
| Mymx | 4.667 | 4.475 | B230312C02Rik | 2.190 | 2.233 | Gm48719 | 1.690 | 3.100 | Stra6l | 3.159 | 1.530 |
| Ripply1 | 4.649 | 9.986 | A330023F24Rik | 2.143 | 2.355 | Gm14319 | 1.689 | 2.575 | Mamdc2 | 3.094 | 2.810 |
| Chrng | 4.631 | 3.816 | Ush1c | 2.141 | 3.097 | Gm16104 | 1.597 | 1.885 | Rgs1 | 2.982 | 2.164 |
| Mymk | 4.619 | 5.664 | Mab21l2 | 2.128 | 2.716 | Fsbp | 1.593 | 3.284 | Dcstamp | 2.979 | 2.685 |
| Cd300c2 | 4.548 | 4.766 | Zbed6 | 2.113 | 3.172 | Mss51 | 1.582 | 3.506 | Rhov | 2.958 | 1.953 |
| Spp1 | 4.487 | 2.205 | Cdh4 | 2.102 | 2.025 | Abca4 | 1.577 | 1.482 | Hk3 | 2.937 | 1.988 |
| Gm12408 | 4.383 | 7.918 | Dchs2 | 2.099 | 2.025 | Gucy2g | 1.563 | 2.180 | Prss46 | 2.936 | 3.120 |
| Rrad | 4.346 | 3.183 | Gm7265 | 2.090 | 2.326 | Zc3h11a | 1.523 | 4.623 | 4930430E12Rik | 2.815 | 2.445 |
| Myh3 | 4.333 | 3.777 | Hoxa11 | 2.089 | 1.694 | Myo1a | 1.514 | 1.424 | Galnt6 | 2.776 | 2.183 |
| Plac8 | 4.303 | 8.288 | A630019I02Rik | 2.083 | 1.810 | Arrdc2 | 1.507 | 1.340 | Lrrc15 | 2.707 | 2.064 |
| Cyfip2 | 4.282 | 5.421 | Mamdc2 | 2.079 | 1.575 | Abcc8 | 1.494 | 4.263 | Fhdc1 | 2.656 | 4.366 |
| Serpina3m | 4.279 | 3.936 | Gm35638 | 2.050 | 3.095 | Usp44 | 1.486 | 2.946 | Pira2 | 2.652 | 2.930 |
| Ccl2 | 4.232 | 5.647 | Amd2 | 2.026 | 3.683 | C1ql3 | 1.482 | 1.397 | Itgb2 | 2.652 | 2.847 |
| Fam167b | 4.211 | 7.291 | Gm49759 | 1.940 | 2.999 | Slc25a25 | 1.476 | 1.906 | Gdf15 | 2.626 | 2.810 |
| Mt3 | 4.191 | 5.334 | Cacna1i | 1.899 | 2.928 | NA | 1.473 | 2.870 | Trem2 | 2.590 | 2.767 |
| Chil3 | 4.189 | 4.593 | NA | 1.882 | 1.886 | Six2 | 1.471 | 2.689 | B430306N03Rik | 2.572 | 2.520 |
| Clec4d | 4.166 | 4.791 | Hoxa13 | 1.839 | 1.582 | Hmga2-ps1 | 1.461 | 1.728 | Pianp | 2.476 | 3.043 |
| Itgax | 4.149 | 4.660 | Tmem178 | 1.814 | 1.594 | Atp13a5 | 1.428 | 2.625 | Dhrs9 | 2.461 | 2.092 |
| Krt18 | 4.101 | 6.708 | 1700001O22Rik | 1.810 | 2.561 | Pfkfb3 | 1.425 | 3.296 | Cd22 | 2.457 | 2.131 |
| Adam8 | 4.092 | 4.067 | Iglon5 | 1.777 | 2.335 | Dpysl5 | 1.425 | 2.948 | Cd180 | 2.433 | 2.767 |
| Mmp3 | 4.072 | 8.766 | Hif3a | 1.746 | 1.728 | F830016B08Rik | 1.424 | 4.031 | Slamf7 | 2.426 | 2.360 |
| Mki67 | 3.990 | 5.109 | Hvcn1 | 1.735 | 1.607 | Atp7a | 1.412 | 2.558 | Tlr8 | 2.410 | 1.556 |
| NA | 3.954 | 5.501 | Col8a2 | 1.722 | 1.721 | NA | 1.402 | 1.766 | Hmmr | 2.387 | 2.062 |

| | | | | | | | | | | | |
|---|---|---|---|---|---|---|---|---|---|---|---|
| Cthrc1 | 3.948 | 5.212 | Spon1 | 1.714 | 2.196 | Arnt2 | 1.400 | 3.021 | Adam8 | 2.386 | 1.969 |
| Gm3636 | 3.944 | 9.846 | Cilp2 | 1.707 | 2.045 | Tph1 | 1.387 | 2.356 | Tlr13 | 2.381 | 2.038 |
| Lgals3 | 3.926 | 5.162 | B430306N03 Rik | 1.674 | 1.478 | Gm34804 | 1.375 | 2.772 | Cd68 | 2.349 | 2.892 |
| Gm45774 | 3.911 | 5.452 | Doc2b | 1.672 | 2.212 | Whrn | 1.347 | 2.712 | Cd300lb | 2.339 | 2.120 |
| Cenpf | 3.904 | 7.785 | Slamf7 | 1.659 | 1.463 | Gm10435 | 1.343 | 2.638 | Ly9 | 2.337 | 2.676 |
| Shisal1 | 3.868 | 8.062 | Pianp | 1.651 | 1.548 | Pde4d | 1.339 | 3.690 | Il1rn | 2.336 | 2.030 |
| Tnnt2 | 3.853 | 5.893 | Tph1 | 1.644 | 3.789 | Adcy9 | 1.337 | 4.541 | Hvcn1 | 2.317 | 2.397 |
| Runx2os1 | 3.841 | 8.841 | Sphkap | 1.621 | 2.547 | Pax3 | 1.336 | 1.705 | Cdca7l | 2.312 | 3.111 |
| Cldn2 | 3.838 | 6.587 | C1ql3 | 1.616 | 1.674 | Ciart | 1.333 | 1.651 | Myo1f | 2.309 | 1.952 |
| Serpinb1a | 3.821 | 9.178 | Gm26604 | 1.613 | 2.046 | Cacna1c | 1.321 | 2.852 | Ctsk | 2.286 | 3.067 |
| Gm49454 | 3.807 | 7.176 | Col19a1 | 1.599 | 2.630 | Tfcp2l1 | 1.298 | 2.577 | A330023F24 Rik | 2.278 | 2.588 |
| Mt2 | 3.751 | 8.615 | Odf3l2 | 1.593 | 1.562 | Prom1 | 1.281 | 3.464 | Gpr137b | 2.267 | 2.597 |
| Ush1c | -2.280 | 2.600 | Adig | -1.818 | 2.764 | Cldn1 | -1.809 | 1.411 | Esrrb | -2.162 | 2.992 |
| Col4a3 | -2.288 | 5.748 | Smtnl1 | -1.824 | 3.333 | Casq2 | -1.814 | 2.498 | Slc7a10 | -2.167 | 1.429 |
| Bcl2 | -2.289 | 6.836 | mt-Tl1 | -1.835 | 2.991 | Tac4 | -1.830 | 1.578 | Fabp3-ps1 | -2.168 | 4.503 |
| Amd1 | -2.307 | 5.843 | Myh7b | -1.844 | 1.541 | Cyfip2 | -1.846 | 1.834 | Apoc1 | -2.173 | 1.859 |
| Efcab6 | -2.327 | 8.537 | Gm28661 | -1.850 | 1.789 | Ssu2 | -1.858 | 1.968 | Cidea | -2.182 | 5.214 |
| Gm8734 | -2.327 | 5.591 | Cidea | -1.858 | 3.901 | Fcna | -1.872 | 1.826 | Paqr9 | -2.184 | 5.184 |
| Prima1 | -2.383 | 4.926 | Gm43743 | -1.858 | 3.605 | Wfdc17 | -1.876 | 3.246 | Eps8l1 | -2.195 | 2.034 |
| Ighm | -2.390 | 5.204 | Zic1 | -1.861 | 3.180 | NA | -1.877 | 3.649 | Klhdc7a | -2.235 | 1.466 |
| Mstn | -2.391 | 6.794 | Myh13 | -1.873 | 1.524 | Gm10925 | -1.878 | 2.238 | Nat8l | -2.287 | 1.526 |
| Srcin1 | -2.400 | 5.406 | Ldhb-ps | -1.904 | 1.768 | Prc1 | -1.884 | 1.463 | Plin5 | -2.399 | 5.078 |
| Tph1 | -2.426 | 3.832 | Ccl7 | -1.916 | 1.913 | 2310043L19R ik | -1.893 | 2.087 | Gm28437 | -2.473 | 2.305 |
| Hipk2 | -2.490 | 6.249 | Plin5 | -1.947 | 3.889 | Hpdl | -1.896 | 3.501 | Hp | -2.486 | 1.561 |
| Cacng7 | -2.499 | 10.07 2 | Acvr1c | -1.952 | 1.545 | Gm41757 | -1.898 | 3.277 | Mb | -2.487 | 2.726 |
| Cd28 | -2.513 | 4.306 | Ch25h | -1.959 | 3.196 | Itgae | -1.913 | 2.233 | Mrap | -2.570 | 1.306 |
| Nat1 | -2.515 | 3.971 | NA | -1.974 | 1.453 | Knstrn | -1.913 | 1.930 | Ces1d | -2.581 | 2.135 |
| Gm33543 | -2.515 | 3.769 | Zdhhc23 | -1.978 | 2.988 | Esrrb | -1.922 | 2.259 | Acsm3 | -2.588 | 3.665 |
| Col9a1 | -2.577 | 4.898 | Gm41757 | -1.980 | 4.778 | Slc25a34 | -1.937 | 1.866 | Scd1 | -2.597 | 3.504 |
| mt-Atp8 | -2.590 | 1.426 | Csrp3 | -1.985 | 1.801 | Cidea | -1.944 | 4.012 | Aspg | -2.597 | 5.689 |
| mt-Nd4l | -2.590 | 1.947 | Wfdc21 | -2.006 | 1.464 | Aspg | -1.954 | 2.670 | Myoz2 | -2.617 | 2.781 |
| Zbed6 | -2.591 | 3.633 | 1700016C15R ik | -2.049 | 2.334 | Gm49540 | -1.992 | 2.558 | mt-Atp6 | -2.706 | 3.590 |
| Hlf | -2.627 | 6.883 | Gm15543 | -2.060 | 3.338 | Gm28437 | -1.998 | 1.389 | Klhl34 | -2.708 | 2.831 |
| Epha7 | -2.632 | 6.355 | Dgat2 | -2.074 | 3.496 | Tnni1 | -2.047 | 1.358 | Arxes2 | -2.776 | 1.490 |
| Cracd | -2.632 | 7.885 | Myoz2 | -2.130 | 2.839 | Gm50221 | -2.054 | 4.480 | Slc36a2 | -2.786 | 3.878 |
| Amd-ps3 | -2.640 | 5.212 | Cytl1 | -2.163 | 1.875 | Dgat2 | -2.063 | 5.314 | Gm49540 | -2.794 | 3.612 |
| Itga4 | -2.679 | 6.834 | Sprr1a | -2.171 | 2.375 | Myom3 | -2.086 | 1.739 | Gm15543 | -2.825 | 5.030 |
| Dusp26 | -2.764 | 7.474 | S100a9 | -2.183 | 4.244 | Gm15543 | -2.136 | 3.702 | Gm29216 | -2.866 | 3.796 |
| Slc15a5 | -2.800 | 3.037 | Klhl34 | -2.184 | 2.642 | Fam240a | -2.151 | 1.599 | Ldhb | -2.866 | 3.293 |
| Smim35 | -2.918 | 7.363 | Ldhb | -2.185 | 2.962 | Gm29216 | -2.156 | 2.023 | Cyp2e1 | -2.866 | 1.546 |
| Abca4 | -2.948 | 3.167 | Lman1l | -2.185 | 2.816 | G0s2 | -2.165 | 4.461 | Mup22 | -2.907 | 1.804 |
| 0610040J01R ik | -3.020 | 7.275 | Gm28437 | -2.255 | 2.283 | BC048679 | -2.193 | 2.156 | Slc25a34 | -2.957 | 3.314 |
| Nt5c1a | -3.023 | 6.266 | Ccl2 | -2.264 | 2.446 | Kcne1l | -2.201 | 1.942 | Fam240a | -2.975 | 2.432 |

| | | | | | | | | | | | |
|---|---|---|---|---|---|---|---|---|---|---|---|
| Helt | -3.026 | 2.970 | Chdh | -2.336 | 1.683 | Klhl34 | -2.222 | 1.864 | Dgat2 | -3.074 | 6.802 |
| Kl | -3.106 | 3.772 | Nalf2 | -2.349 | 4.424 | Ldhb | -2.346 | 2.429 | Cdo1 | -3.125 | 2.092 |
| Kcnab1 | -3.128 | 5.809 | mt-Atp6 | -2.432 | 2.816 | Ccl2 | -2.362 | 3.043 | Cfd | -3.205 | 2.269 |
| Ppp1r1a | -3.135 | 10.693 | Barx2 | -2.490 | 1.982 | Ankrd2 | -2.389 | 2.715 | mt-Ts1 | -3.211 | 2.925 |
| Ace2 | -3.145 | 5.012 | BC048679 | -2.503 | 2.100 | Tnnt1 | -2.444 | 1.529 | Ankrd2 | -3.212 | 3.085 |
| Dach2 | -3.219 | 1.552 | Kcne1l | -2.610 | 3.563 | Lman1l | -2.507 | 2.260 | Adig | -3.331 | 5.990 |
| Tiam1 | -3.254 | 8.097 | Chil3 | -2.646 | 2.595 | Mal | -2.603 | 2.168 | A530016L24Rik | -3.349 | 2.220 |
| Amd-ps4 | -3.262 | 5.072 | Gm50221 | -2.655 | 5.710 | Csrp3 | -2.651 | 1.784 | Adipoq | -3.462 | 1.671 |
| Gm7265 | -3.297 | 5.177 | Ankrd2 | -2.660 | 2.938 | Mpz | -2.716 | 1.830 | Myl10 | -3.467 | 2.137 |
| A630019I02Rik | -3.364 | 5.169 | Slc25a34 | -2.735 | 3.625 | mt-Ts1 | -2.772 | 1.856 | BC048679 | -3.468 | 3.535 |
| Stk26 | -3.404 | 8.578 | mt-Ts1 | -2.759 | 2.825 | H2-Q10 | -2.916 | 2.053 | Plin1 | -3.536 | 1.840 |
| Lamc3 | -3.408 | 4.370 | Myl10 | -3.121 | 2.578 | Myoz2 | -2.920 | 2.636 | Cidec | -3.568 | 1.329 |
| Snph | -3.546 | 9.959 | Gm29216 | -3.132 | 4.501 | Myl10 | -2.925 | 1.502 | Retn | -3.756 | 2.016 |
| Dmd | -3.561 | 11.853 | 9430073C21Rik | -3.618 | 3.325 | Barx2 | -3.031 | 1.701 | 9430073C21Rik | -3.994 | 2.874 |
| Cdh4 | -3.959 | 3.649 | Bdh1 | -3.898 | 2.553 | Chil3 | -3.127 | 3.147 | Pck1 | -4.40 | 1.559 |
| Tafa4 | -4.452 | 8.554 | Myh7 | -3.966 | 1.316 | Ccl7 | -3.332 | 2.681 | Bdh1 | -4.50 | 3.062 |
| 1700001O22Rik | -5.019 | 5.535 | Myh2 | -3.992 | 2.457 | 9430073C21Rik | -3.522 | 1.968 | Myh2 | -5.487 | 3.29 |
| Themis3 | -6.115 | 7.133 | Myl3 | -5.501 | 1.967 | Myh2 | -3.633 | 2.201 | Myl2 | -6.104 | 2.190 |
| Amd2 | -6.245 | 7.586 | Myl2 | -5.749 | 2.131 | Bdh1 | -4.246 | 1.765 | Myl3 | -8.056 | 2.077 |

### Supplementary Table 4. Significantly upregulated and downregulated lipids

| *mdx*/WT | | | *mdx* ASA/VEH | | | *mdx* DMF/VEH | | | *mdx* RIB/VEH | | |
|---|---|---|---|---|---|---|---|---|---|---|---|
| Lipid | log2(FC) | -log$_{10}$(*p*) | Lipid | log2(FC) | -log$_{10}$(*p*) | Lipid | log2(FC) | -log$_{10}$(*p*) | Lipid | log2(FC) | -log$_{10}$(*p*) |
| TG-D5 (IS) | 1.544 | 1.166 | dhCer 20:0 | -1.608 | 1.826 | | | | TG(O-52:2) | 1.003 | 1.428 |
| AcylCarnitine 14:2 | 1.478 | 1.916 | | | | | | | PC(P-36:2) | 1.101 | 1.393 |
| AcylCarnitine 14:1 | 1.421 | 2.508 | | | | | | | TG 18:2_18:2_18:2 | -1.012 | 3.957 |
| AcylCarnitine 12:0 | 1.247 | 2.029 | | | | | | | TG 18:0_18:0_18:0 | -1.051 | 2.955 |
| TG 14:1_16:1_18:0 | 1.179 | 2.586 | | | | | | | DG 30:0 -(14:0) | -1.054 | 2.346 |
| AcylCarnitine 16:1 | 1.144 | 2.116 | | | | | | | TG 18:0_18:0_18:1 | -1.056 | 3.032 |
| TG(O-52:2) | 1.102 | 3.182 | | | | | | | TG 14:0_18:2_18:2 | -1.070 | 2.728 |
| PC(P-32:1) | 1.092 | 4.471 | | | | | | | DG 32:0 -(16:0) | -1.129 | 3.814 |
| PC(O-32:2) | 1.058 | 5.022 | | | | | | | | | |
| PE(P-20:0_22:6) | 1.045 | 6.511 | | | | | | | | | |
| PC(P-34:2) | 1.023 | 6.434 | | | | | | | | | |
| PI 40:5 | 1.020 | 2.500 | | | | | | | | | |
| Hex2Cer(d18:1_18:0) | 1.018 | 1.910 | | | | | | | | | |
| PI 40:6 | 1.017 | 2.123 | | | | | | | | | |
| SM 37:2 | 1.010 | 5.991 | | | | | | | | | |
| SM 34:2 | -1.009 | 5.908 | | | | | | | | | |
| PE(P-16:0_18:3) | -1.028 | 5.711 | | | | | | | | | |
| TG 18:0_18:0_18:0 | -1.038 | 3.036 | | | | | | | | | |
| PI 36:1 | -1.082 | 2.017 | | | | | | | | | |
| Hex3Cer(d18:1_22:0) | -1.086 | 2.531 | | | | | | | | | |
| PC 36:1 | -1.097 | 4.093 | | | | | | | | | |
| PE 34:1 | -1.112 | 6.763 | | | | | | | | | |
| DG 36:4 -(18:2) | -1.112 | 1.619 | | | | | | | | | |
| PC(P-36:3) | -1.154 | 4.996 | | | | | | | | | |
| SM 34:0 | -1.161 | 7.164 | | | | | | | | | |
| PC(O-36:4) | -1.162 | 5.050 | | | | | | | | | |
| PE 17:0/17:0 (IS) | -1.196 | 6.773 | | | | | | | | | |
| PE(P-16:0_18:1) | -1.200 | 5.098 | | | | | | | | | |
| LPC(O-16:0) | -1.313 | 6.418 | | | | | | | | | |
| SM 40:0 | -1.331 | 4.162 | | | | | | | | | |
| LPC 18:3(104) | -1.369 | 5.545 | | | | | | | | | |
| LPC(O-18:1) | -1.369 | 3.758 | | | | | | | | | |
| PC 40:4 | -1.390 | 4.896 | | | | | | | | | |
| SM 44:3 | -1.397 | 5.464 | | | | | | | | | |
| LPC(P-18:0) | -1.430 | 3.893 | | | | | | | | | |
| PC(P-40:4) | -1.470 | 3.679 | | | | | | | | | |

| | | |
|---|---|---|
| Hex3Cer(d18:1_16:0) | -1.489 | 3.890 |
| PC(O-40:5) | -1.565 | 5.289 |
| Hex2Cer(d18:1_22:0) | -1.667 | 4.306 |
| Hex2Cer(d18:1_24:0) | -1.778 | 5.547 |
| PE 35:1 | -1.801 | 6.465 |
| PE 40:4 | -1.829 | 6.929 |
| Hex2Cer(d18:1_24:1) | -2.155 | 5.269 |
| Hex1Cer(d18:1_16:0) | -2.320 | 4.957 |
| CE 22:1 | -2.387 | 2.392 |
| PE 36:1 | -2.470 | 7.661 |
| PE 36:0 | -2.478 | 7.519 |
| CE 20:1 | -2.486 | 1.593 |
| CE 20:2 | -2.655 | 1.407 |
| GM3(d18:1_16:0) | -2.685 | 6.432 |
| Hex2Cer(d18:1_16:0) | -2.929 | 7.057 |
| CE 22:4 | -3.740 | 2.231 |
| CE 24:1 | -3.741 | 5.136 |
| CE 24:4 | -5.042 | 3.830 |

#### Supplementary Table 5. Metabolites Pathway Analysis *mdx* ASA/VEH

| Pathway | Total metabolites in pathway | Hits | Raw *p* value | -log$_{10}$(*p*) | Impact |
|---|---|---|---|---|---|
| **Significant Pathways** | | | | | |
| Butyrate Metabolism | 16 | 5 | 3.9E-06 | 5.406 | 0.359 |
| Phytanic Acid Peroxisomal Oxidation | 19 | 5 | 1.0E-05 | 4.995 | 0.000 |
| Glycerolipid Metabolism | 23 | 5 | 2.8E-05 | 4.553 | 0.390 |
| Ketone Body Metabolism | 12 | 4 | 3.3E-05 | 4.485 | 0.046 |
| Citric Acid Cycle | 25 | 5 | 4.3E-05 | 4.364 | 0.531 |
| Threonine and 2-Oxobutanoate Degradation | 13 | 4 | 4.7E-05 | 4.330 | 0.143 |
| Glutamate Metabolism | 45 | 6 | 7.1E-05 | 4.147 | 0.172 |
| Ethanol Degradation | 15 | 4 | 8.7E-05 | 4.059 | 0.038 |
| Mitochondrial Electron Transport Chain | 15 | 4 | 8.7E-05 | 4.059 | 0.286 |
| Galactose Metabolism | 31 | 5 | 1.3E-04 | 3.888 | 0.009 |
| Mitochondrial Beta-Oxidation of Short Chain Saturated Fatty Acids | 17 | 4 | 1.5E-04 | 3.827 | 0.630 |
| Transfer of Acetyl Groups into Mitochondria | 18 | 4 | 1.9E-04 | 3.723 | 0.127 |
| Propanoate Metabolism | 36 | 5 | 2.7E-04 | 3.566 | 0.458 |
| De Novo Triacylglycerol Biosynthesis | 9 | 3 | 3.8E-04 | 3.421 | 0.200 |
| Mitochondrial Beta-Oxidation of Medium Chain Saturated Fatty Acids | 22 | 4 | 4.3E-04 | 3.364 | 0.650 |
| Mitochondrial Beta-Oxidation of Long Chain Saturated Fatty Acids | 24 | 4 | 6.1E-04 | 3.212 | 0.422 |
| Cardiolipin Biosynthesis | 11 | 3 | 7.3E-04 | 3.137 | 0.299 |
| Beta Oxidation of Very Long Chain Fatty Acids | 13 | 3 | 1.2E-03 | 2.908 | 0.544 |
| Valine, Leucine and Isoleucine Degradation | 51 | 5 | 1.4E-03 | 2.844 | 0.246 |
| Pyruvate Metabolism | 30 | 4 | 1.5E-03 | 2.830 | 0.434 |
| Amino Sugar Metabolism | 31 | 4 | 1.7E-03 | 2.775 | 0.000 |
| Fatty Acid Metabolism | 40 | 4 | 4.4E-03 | 2.356 | 0.688 |
| Lysine Degradation | 20 | 3 | 4.6E-03 | 2.342 | 0.093 |
| Caffeine Metabolism | 20 | 3 | 4.6E-03 | 2.342 | 0.250 |
| Phenylacetate Metabolism | 8 | 2 | 8.0E-03 | 2.099 | 0.500 |
| Glycerol Phosphate Shuttle | 8 | 2 | 8.0E-03 | 2.099 | 0.762 |
| Starch and Sucrose Metabolism | 26 | 3 | 9.7E-03 | 2.013 | 0.000 |
| Beta-Alanine Metabolism | 26 | 3 | 9.7E-03 | 2.013 | 0.190 |
| Glycine and Serine Metabolism | 50 | 4 | 9.9E-03 | 2.004 | 0.104 |
| Pentose Phosphate Pathway | 27 | 3 | 1.1E-02 | 1.967 | 0.154 |
| Tryptophan Metabolism | 55 | 4 | 1.4E-02 | 1.859 | 0.004 |
| Retinol Metabolism | 30 | 3 | 1.4E-02 | 1.839 | 0.082 |
| Bile Acid Biosynthesis | 60 | 4 | 1.9E-02 | 1.728 | 0.111 |
| Fatty Acid Elongation In Mitochondria | 33 | 3 | 1.9E-02 | 1.726 | 0.411 |
| Purine Metabolism | 63 | 4 | 2.2E-02 | 1.656 | 0.129 |
| Lactose Synthesis | 14 | 2 | 2.4E-02 | 1.615 | 0.221 |

| | | | | | |
|---|---|---|---|---|---|
| Methionine Metabolism | 39 | 3 | 2.9E-02 | 1.531 | 0.000 |
| Carnitine Synthesis | 16 | 2 | 3.1E-02 | 1.504 | 0.000 |
| Nucleotide Sugars Metabolism | 16 | 2 | 3.1E-02 | 1.504 | 0.000 |
| Steroid Biosynthesis | 43 | 3 | 3.8E-02 | 1.420 | 0.000 |
| Betaine Metabolism | 18 | 2 | 3.9E-02 | 1.407 | 0.000 |
| Pantothenate and CoA Biosynthesis | 19 | 2 | 4.3E-02 | 1.364 | 0.000 |
| Glycolysis | 20 | 2 | 4.8E-02 | 1.323 | 0.134 |
| **Non-Significant Pathways** | | | | | |
| Arginine and Proline Metabolism | 48 | 3 | 5.0E-02 | 1.298 | 0.097 |
| Urea Cycle | 23 | 2 | 6.1E-02 | 1.212 | 0.000 |
| Folate Metabolism | 24 | 2 | 6.6E-02 | 1.179 | 0.005 |
| Cysteine Metabolism | 24 | 2 | 6.6E-02 | 1.179 | 0.090 |
| Phospholipid Biosynthesis | 25 | 2 | 7.1E-02 | 1.147 | 0.113 |
| Ammonia Recycling | 25 | 2 | 7.1E-02 | 1.147 | 0.145 |
| Selenoamino Acid Metabolism | 28 | 2 | 8.7E-02 | 1.060 | 0.000 |
| Fructose and Mannose Degradation | 28 | 2 | 8.7E-02 | 1.060 | 0.000 |
| Gluconeogenesis | 30 | 2 | 9.8E-02 | 1.009 | 0.112 |
| Nicotinate and Nicotinamide Metabolism | 32 | 2 | 1.1E-01 | 0.961 | 0.249 |
| Fatty Acid Biosynthesis | 33 | 2 | 1.2E-01 | 0.938 | 0.000 |
| Biotin Metabolism | 7 | 1 | 1.2E-01 | 0.924 | 0.000 |
| Homocysteine Degradation | 7 | 1 | 1.2E-01 | 0.924 | 0.000 |
| Malate-Aspartate Shuttle | 7 | 1 | 1.2E-01 | 0.924 | 0.000 |
| Aspartate Metabolism | 34 | 2 | 1.2E-01 | 0.916 | 0.000 |
| Histidine Metabolism | 35 | 2 | 1.3E-01 | 0.895 | 0.000 |
| Glucose-Alanine Cycle | 9 | 1 | 1.5E-01 | 0.822 | 0.000 |
| Lactose Degradation | 9 | 1 | 1.5E-01 | 0.822 | 0.000 |
| Thiamine Metabolism | 9 | 1 | 1.5E-01 | 0.822 | 0.000 |
| Trehalose Degradation | 11 | 1 | 1.8E-01 | 0.742 | 0.000 |
| Phosphatidylethanolamine Biosynthesis | 13 | 1 | 2.1E-01 | 0.677 | 0.000 |
| Alanine Metabolism | 14 | 1 | 2.2E-01 | 0.648 | 0.000 |
| Phosphatidylinositol Phosphate Metabolism | 14 | 1 | 2.2E-01 | 0.648 | 0.000 |
| Riboflavin Metabolism | 14 | 1 | 2.2E-01 | 0.648 | 0.000 |
| Spermidine and Spermine Biosynthesis | 14 | 1 | 2.2E-01 | 0.648 | 0.000 |
| Pyrimidine Metabolism | 54 | 2 | 2.5E-01 | 0.600 | 0.216 |
| Plasmalogen Synthesis | 16 | 1 | 2.5E-01 | 0.597 | 0.000 |
| Pterine Biosynthesis | 18 | 1 | 2.8E-01 | 0.553 | 0.000 |
| Phosphatidylcholine Biosynthesis | 18 | 1 | 2.8E-01 | 0.553 | 0.098 |
| Sulfate/Sulfite Metabolism | 19 | 1 | 2.9E-01 | 0.533 | 0.000 |
| Glutathione Metabolism | 19 | 1 | 2.9E-01 | 0.533 | 0.168 |
| Estrone Metabolism | 20 | 1 | 3.1E-01 | 0.515 | 0.160 |
| Inositol Phosphate Metabolism | 22 | 1 | 3.3E-01 | 0.480 | 0.226 |
| Androstenedione Metabolism | 23 | 1 | 3.4E-01 | 0.464 | 0.161 |

| | | | | | |
|---|---|---|---|---|---|
| Inositol Metabolism | 28 | 1 | 4.0E-01 | 0.397 | 0.140 |
| Androgen and Estrogen Metabolism | 29 | 1 | 4.1E-01 | 0.385 | 0.104 |
| Porphyrin Metabolism | 36 | 1 | 4.8E-01 | 0.315 | 0.000 |
| Sphingolipid Metabolism | 36 | 1 | 4.8E-01 | 0.315 | 0.028 |
| Steroidogenesis | 42 | 1 | 5.4E-01 | 0.268 | 0.024 |
| Tyrosine Metabolism | 55 | 1 | 6.4E-01 | 0.194 | 0.097 |

**Supplementary Table 6. Metabolites Pathway Analysis *mdx* DMF/VEH**

| Pathway | Total metabolites in pathways | Hits | Raw *p* value | $-\log_{10}(p)$ | Impact |
|---|---|---|---|---|---|
| **Significant Pathways** | | | | | |
| Pentose Phosphate Pathway | 27 | 3 | 6.81E-05 | 4.166 | 0.1926 |
| Glycolysis | 20 | 2 | 0.002 | 2.656 | 0.034 |
| Gluconeogenesis | 30 | 2 | 0.004 | 2.302 | 0.058 |
| **Non-Significant Pathways** | | | | | |
| Nucleotide Sugars Metabolism | 16 | 1 | 0.062 | 1.205 | 0 |
| Inositol Phosphate Metabolism | 22 | 1 | 0.084 | 1.071 | 0 |
| Starch and Sucrose Metabolism | 26 | 1 | 0.099 | 1.001 | 0 |
| Inositol Metabolism | 28 | 1 | 0.107 | 0.971 | 0 |
| Fructose and Mannose Degradation | 28 | 1 | 0.107 | 0.971 | 0.203 |
| Galactose Metabolism | 31 | 1 | 0.117 | 0.928 | 0 |
| Amino Sugar Metabolism | 31 | 1 | 0.117 | 0.928 | 0.041 |
| Glutamate Metabolism | 45 | 1 | 0.167 | 0.775 | 0 |
| Purine Metabolism | 63 | 1 | 0.228 | 0.641 | 0.022 |

#### Supplementary Table 7. Metabolites Pathway Analysis *mdx* RIB/VEH

| Pathway | Total metabolites in pathway | Hits | Raw *p* value | -$\log_{10}(p)$ | Impact |
|---|---|---|---|---|---|
| **Significant Pathway** | | | | | |
| Galactose metabolism | 27 | 2 | 0.014 | 1.832 | 0.090 |
| **Non-Significant Pathways** | | | | | |
| Taurine and hypotaurine metabolism | 8 | 1 | 0.055 | 1.254 | 0 |
| Arginine biosynthesis | 14 | 1 | 0.095 | 1.019 | 0 |
| Butanoate metabolism | 15 | 1 | 0.102 | 0.990 | 0 |
| Starch and sucrose metabolism | 15 | 1 | 0.102 | 0.990 | 0.087 |
| Pentose and glucuronate interconversions | 19 | 1 | 0.127 | 0.893 | 0.168 |
| Citrate cycle (TCA cycle) | 20 | 1 | 0.134 | 0.872 | 0.032 |
| Pantothenate and CoA biosynthesis | 20 | 1 | 0.134 | 0.872 | 0.166 |
| Propanoate metabolism | 22 | 1 | 0.146 | 0.834 | 0 |
| Pentose phosphate pathway | 23 | 1 | 0.152 | 0.816 | 0.004 |
| Alanine, aspartate and glutamate metabolism | 28 | 1 | 0.182 | 0.737 | 0 |
| Pyrimidine metabolism | 39 | 1 | 0.245 | 0.609 | 0.046 |
| Fatty acid degradation | 39 | 1 | 0.245 | 0.609 | 0.124 |
| Primary bile acid biosynthesis | 46 | 1 | 0.283 | 0.546 | 0.022 |
| Steroid hormone biosynthesis | 79 | 1 | 0.439 | 0.356 | 0.013 |

#### Supplementary Table 8. Integrative Analysis of Significant Genes & Metabolites *mdx*/WT

| Pathway | Total metabolites & genes in pathway | Metabolite hits | Gene hits | Raw *p* value | $-\log_{10}(p)$ | Impact |
|---|---|---|---|---|---|---|
| **Pathways with Gene and Metabolite Hits and Impact Score >0.2** | | | | | | |
| Purine metabolism | 173 | 2 | 13 | 0.394 | 0.405 | 0.576 |
| Glycine, serine and threonine metabolism | 72 | 1 | 5 | 0.426 | 0.370 | 0.563 |
| Pyrimidine metabolism | 100 | 1 | 8 | 0.492 | 0.308 | 0.535 |
| Arginine and proline metabolism | 74 | 2 | 11 | 0.002 | 2.725 | 0.521 |
| Glycerophospholipid metabolism | 87 | 2 | 10 | 0.032 | 1.497 | 0.500 |
| Galactose metabolism | 51 | 2 | 6 | 0.010 | 2.013 | 0.480 |
| Glutathione metabolism | 57 | 1 | 6 | 0.129 | 0.888 | 0.393 |
| Histidine metabolism | 32 | 1 | 4 | 0.072 | 1.140 | 0.387 |
| Ether lipid metabolism | 40 | 1 | 7 | 0.012 | 1.917 | 0.385 |
| Sphingolipid metabolism | 81 | 1 | 7 | 0.378 | 0.423 | 0.350 |
| Tryptophan metabolism | 84 | 2 | 3 | 0.528 | 0.277 | 0.241 |
| beta-Alanine metabolism | 44 | 1 | 3 | 0.354 | 0.452 | 0.233 |
| One carbon pool by folate | 72 | 1 | 1 | 0.990 | 0.004 | 0.225 |
| **Pathways with Gene and Metabolite Hits and Impact Score <0.2** | | | | | | |
| Alanine, aspartate and glutamate metabolism | 61 | 1 | 3 | 0.628 | 0.202 | 0.183 |
| Lipoic acid metabolism | 51 | 1 | 1 | 0.726 | 0.139 | 0.160 |
| Glyoxylate and dicarboxylate metabolism | 56 | 1 | 1 | 0.735 | 0.134 | 0.091 |
| Glycosylphosphatidylinositol (GPI)-anchor biosynthesis | 57 | 1 | 2 | 0.784 | 0.106 | 0.089 |
| Aminoacyl-tRNA biosynthesis | 74 | 3 | 1 | 0.128 | 0.893 | 0.068 |
| **Other Pathways** | | | | | | |
| Neomycin, kanamycin and gentamicin biosynthesis | 4 | 0 | 2 | 0.019 | 1.726 | 1.333 |
| Valine, leucine and isoleucine biosynthesis | 12 | 0 | 2 | 0.093 | 1.030 | 0.909 |
| Nicotinate and nicotinamide metabolism | 43 | 0 | 8 | 0.029 | 1.540 | 0.667 |
| Glycosaminoglycan biosynthesis - chondroitin sulfate or dermatan sulfate | 18 | 0 | 4 | 0.015 | 1.814 | 0.588 |
| Fructose and mannose metabolism | 37 | 0 | 3 | 0.498 | 0.303 | 0.500 |
| Glycerolipid metabolism | 34 | 0 | 5 | 0.088 | 1.054 | 0.455 |
| Pyruvate metabolism | 48 | 0 | 6 | 0.116 | 0.935 | 0.426 |
| Glycolysis or Gluconeogenesis | 60 | 0 | 6 | 0.322 | 0.493 | 0.407 |
| Amino sugar and nucleotide sugar metabolism | 93 | 0 | 8 | 0.403 | 0.394 | 0.391 |
| Tyrosine metabolism | 88 | 0 | 4 | 0.900 | 0.046 | 0.391 |
| Glycosphingolipid biosynthesis - ganglio series | 47 | 0 | 3 | 0.718 | 0.144 | 0.348 |
| Ascorbate and aldarate metabolism | 19 | 0 | 2 | 0.357 | 0.447 | 0.333 |
| Pentose and glucuronate interconversions | 34 | 0 | 2 | 0.634 | 0.198 | 0.303 |
| Phenylalanine metabolism | 21 | 0 | 2 | 0.460 | 0.337 | 0.300 |
| Citrate cycle (TCA cycle) | 42 | 0 | 3 | 0.602 | 0.220 | 0.268 |

| | | | | | | |
|---|---|---|---|---|---|---|
| Riboflavin metabolism | 9 | 0 | 1 | 0.524 | 0.281 | 0.250 |
| Metabolism of xenobiotics by cytochrome P450 | 118 | 0 | 2 | 0.997 | 0.001 | 0.239 |
| Glycosaminoglycan degradation | 44 | 0 | 3 | 0.569 | 0.245 | 0.233 |
| Cysteine and methionine metabolism | 72 | 0 | 6 | 0.454 | 0.343 | 0.225 |
| Starch and sucrose metabolism | 37 | 0 | 3 | 0.602 | 0.220 | 0.222 |
| Nitrogen metabolism | 10 | 0 | 2 | 0.093 | 1.030 | 0.222 |
| Inositol phosphate metabolism | 69 | 0 | 5 | 0.642 | 0.192 | 0.206 |
| Vitamin B6 metabolism | 21 | 0 | 1 | 0.833 | 0.080 | 0.200 |
| Mucin type O-glycan biosynthesis | 27 | 0 | 2 | 0.553 | 0.257 | 0.192 |
| Arachidonic acid metabolism | 93 | 0 | 5 | 0.843 | 0.074 | 0.174 |
| Valine, leucine and isoleucine degradation | 87 | 0 | 3 | 0.975 | 0.011 | 0.174 |
| Pantothenate and CoA biosynthesis | 36 | 0 | 2 | 0.670 | 0.174 | 0.171 |
| Drug metabolism - cytochrome P450 | 39 | 0 | 2 | 0.733 | 0.135 | 0.158 |
| Arginine biosynthesis | 28 | 0 | 1 | 0.876 | 0.057 | 0.148 |
| Glycosaminoglycan biosynthesis - heparan sulfate or heparin | 15 | 0 | 1 | 0.647 | 0.189 | 0.143 |
| Folate biosynthesis | 60 | 0 | 3 | 0.856 | 0.068 | 0.136 |
| Linoleic acid metabolism | 17 | 0 | 1 | 0.833 | 0.080 | 0.125 |
| Caffeine metabolism | 21 | 0 | 1 | 0.738 | 0.132 | 0.100 |
| Fatty acid degradation | 102 | 0 | 3 | 0.996 | 0.002 | 0.099 |
| alpha-Linolenic acid metabolism | 22 | 0 | 1 | 0.738 | 0.132 | 0.095 |
| D-Amino acid metabolism | 23 | 0 | 1 | 0.696 | 0.158 | 0.091 |
| Retinol metabolism | 45 | 0 | 1 | 0.989 | 0.005 | 0.091 |
| Propanoate metabolism | 46 | 0 | 2 | 0.865 | 0.063 | 0.089 |
| Pentose phosphate pathway | 49 | 0 | 1 | 0.980 | 0.009 | 0.083 |
| Fatty acid biosynthesis | 129 | 0 | 2 | 1.000 | 0.000 | 0.078 |
| Steroid biosynthesis | 81 | 0 | 2 | 0.982 | 0.008 | 0.075 |
| Lysine degradation | 57 | 0 | 3 | 0.742 | 0.129 | 0.071 |
| Steroid hormone biosynthesis | 177 | 0 | 2 | 1.000 | 0.000 | 0.068 |
| Selenocompound metabolism | 35 | 0 | 1 | 0.893 | 0.049 | 0.059 |
| Glycosphingolipid biosynthesis - globo and isoglobo series | 37 | 0 | 1 | 0.942 | 0.026 | 0.056 |
| Glycosphingolipid biosynthesis - lacto and neolacto series | 104 | 0 | 2 | 0.999 | 0.000 | 0.039 |
| Fatty acid elongation | 75 | 0 | 1 | 0.997 | 0.001 | 0.027 |
| N-Glycan biosynthesis | 81 | 0 | 1 | 0.998 | 0.001 | 0.025 |
| Primary bile acid biosynthesis | 90 | 1 | 0 | 0.347 | 0.459 | 0.022 |
| Porphyrin metabolism | 55 | 1 | 0 | 0.249 | 0.604 | 0.019 |

#### Supplementary Table 9. Integrative Analysis of Significant Genes & Metabolites *mdx* ASA/VEH

| Pathway | Total metabolites & genes in pathway | Metabolite hits | Gene hits | Raw *p* value | $-\log_{10}(p)$ | Impact |
|---|---|---|---|---|---|---|
| **Pathways with Gene and Metabolite Hits and Impact Score >0.2** | | | | | | |
| Glycerolipid metabolism | 34 | 2 | 2 | 0.009 | 2.042 | 0.424 |
| Butanoate metabolism | 29 | 2 | 2 | 0.004 | 2.374 | 0.393 |
| Propanoate metabolism | 46 | 3 | 1 | 0.031 | 1.513 | 0.311 |
| Pyrimidine metabolism | 100 | 1 | 3 | 0.283 | 0.549 | 0.303 |
| Purine metabolism | 173 | 3 | 2 | 0.476 | 0.322 | 0.273 |
| Pyruvate metabolism | 48 | 1 | 1 | 0.361 | 0.443 | 0.234 |
| **Pathways with Gene and Metabolite Hits and Impact Score <0.2** | | | | | | |
| Glutathione metabolism | 57 | 1 | 2 | 0.183 | 0.736 | 0.179 |
| Cysteine and methionine metabolism | 72 | 1 | 4 | 0.046 | 1.342 | 0.169 |
| Glycolysis or Gluconeogenesis | 60 | 1 | 1 | 0.498 | 0.303 | 0.119 |
| beta-Alanine metabolism | 44 | 1 | 1 | 0.326 | 0.487 | 0.116 |
| Glycerophospholipid metabolism | 87 | 1 | 1 | 0.703 | 0.153 | 0.116 |
| Fatty acid elongation | 75 | 1 | 1 | 0.576 | 0.240 | 0.108 |
| Tryptophan metabolism | 84 | 1 | 3 | 0.179 | 0.747 | 0.084 |
| Valine, leucine and isoleucine degradation | 87 | 1 | 1 | 0.685 | 0.164 | 0.081 |
| Inositol phosphate metabolism | 69 | 1 | 1 | 0.568 | 0.246 | 0.074 |
| Alanine, aspartate and glutamate metabolism | 61 | 1 | 1 | 0.486 | 0.313 | 0.050 |
| Glycine, serine and threonine metabolism | 72 | 1 | 1 | 0.560 | 0.252 | 0.042 |
| One carbon pool by folate | 72 | 1 | 1 | 0.653 | 0.185 | 0.042 |
| Primary bile acid biosynthesis | 90 | 1 | 1 | 0.648 | 0.188 | 0.034 |
| **Other Pathways** | | | | | | |
| Biosynthesis of unsaturated fatty acids | 47 | 0 | 1 | 0.306 | 0.514 | 0.478 |
| Phenylalanine, tyrosine and tryptophan biosynthesis | 11 | 0 | 1 | 0.207 | 0.684 | 0.400 |
| Metabolism of xenobiotics by cytochrome P450 | 118 | 0 | 2 | 0.534 | 0.272 | 0.342 |
| Fatty acid degradation | 102 | 2 | 0 | 0.074 | 1.130 | 0.317 |
| Citrate cycle (TCA cycle) | 42 | 2 | 0 | 0.022 | 1.664 | 0.293 |
| Nitrogen metabolism | 10 | 0 | 1 | 0.124 | 0.907 | 0.222 |
| Pentose phosphate pathway | 49 | 2 | 0 | 0.028 | 1.549 | 0.208 |
| Glycosaminoglycan biosynthesis - heparan sulfate or heparin | 15 | 0 | 1 | 0.207 | 0.684 | 0.143 |
| Drug metabolism - other enzymes | 69 | 0 | 4 | 0.034 | 1.472 | 0.132 |
| Pentose and glucuronate interconversions | 34 | 1 | 0 | 0.201 | 0.696 | 0.121 |
| Glycosaminoglycan biosynthesis - chondroitin sulfate or dermatan sulfate | 18 | 0 | 1 | 0.233 | 0.633 | 0.118 |
| Arachidonic acid metabolism | 93 | 0 | 2 | 0.492 | 0.308 | 0.109 |
| Phenylalanine metabolism | 21 | 0 | 1 | 0.306 | 0.514 | 0.100 |

| | | | | | | |
|---|---|---|---|---|---|---|
| Galactose metabolism | 51 | 3 | 0 | 0.003 | 2.482 | 0.100 |
| Nicotinate and nicotinamide metabolism | 43 | 0 | 1 | 0.609 | 0.215 | 0.095 |
| Retinol metabolism | 45 | 0 | 1 | 0.635 | 0.197 | 0.091 |
| Valine, leucine and isoleucine biosynthesis | 12 | 1 | 0 | 0.090 | 1.046 | 0.091 |
| Terpenoid backbone biosynthesis | 36 | 1 | 0 | 0.192 | 0.717 | 0.086 |
| Starch and sucrose metabolism | 37 | 1 | 0 | 0.162 | 0.789 | 0.083 |
| Arginine and proline metabolism | 74 | 0 | 4 | 0.031 | 1.509 | 0.082 |
| Fatty acid biosynthesis | 129 | 1 | 0 | 0.423 | 0.374 | 0.078 |
| Glyoxylate and dicarboxylate metabolism | 56 | 1 | 0 | 0.316 | 0.500 | 0.073 |
| Taurine and hypotaurine metabolism | 17 | 1 | 0 | 0.090 | 1.046 | 0.063 |
| Pantothenate and CoA biosynthesis | 36 | 1 | 0 | 0.211 | 0.676 | 0.057 |
| Drug metabolism - cytochrome P450 | 39 | 0 | 1 | 0.451 | 0.345 | 0.053 |
| Arginine biosynthesis | 28 | 1 | 0 | 0.152 | 0.817 | 0.037 |
| Folate biosynthesis | 60 | 0 | 1 | 0.670 | 0.174 | 0.034 |
| Steroid biosynthesis | 81 | 0 | 1 | 0.741 | 0.130 | 0.025 |
| Tyrosine metabolism | 88 | 0 | 1 | 0.790 | 0.103 | 0.023 |
| Lipoic acid metabolism | 51 | 1 | 0 | 0.283 | 0.549 | 0.020 |
| Glycosphingolipid biosynthesis - lacto and neolacto series | 104 | 0 | 1 | 0.871 | 0.060 | 0.019 |
| Lysine degradation | 57 | 1 | 0 | 0.300 | 0.523 | 0.018 |
| Steroid hormone biosynthesis | 177 | 0 | 1 | 0.967 | 0.014 | 0.011 |
| Amino sugar and nucleotide sugar metabolism | 93 | 1 | 0 | 0.394 | 0.405 | 0.011 |

**Supplementary Table 10. Integrative Analysis of Significant Genes & Metabolites *mdx* DMF/VEH.**

| Pathway | Total metabolites & genes in pathway | Metabolite hits | Gene hits | Raw *p* value | $-\log_{10}(p)$ | Impact |
|---|---|---|---|---|---|---|
| **Pathway with Gene and Metabolite Hits and Impact Score >0.2** | | | | | | |
| Fructose and mannose metabolism | 37 | 1 | 2 | 0.041 | 1.386 | 0.500 |
| Pentose phosphate pathway | 49 | 2 | 1 | 0.056 | 1.256 | 0.333 |
| Purine metabolism | 173 | 1 | 5 | 0.297 | 0.527 | 0.308 |
| Glycolysis or Gluconeogenesis | 60 | 1 | 3 | 0.056 | 1.252 | 0.305 |
| Amino sugar and nucleotide sugar metabolism | 93 | 1 | 2 | 0.339 | 0.470 | 0.239 |
| **Pathways with Gene and Metabolite Hits and Impact Score <0.2** | | | | | | |
| Inositol phosphate metabolism | 69 | 1 | 1 | 0.470 | 0.327 | 0.074 |
| **Other Pathways** | | | | | | |
| Neomycin, kanamycin and gentamicin biosynthesis | 4 | 1 | 0 | 0.005 | 2.285 | 0.667 |
| Pyrimidine metabolism | 100 | 0 | 4 | 0.299 | 0.524 | 0.444 |
| Starch and sucrose metabolism | 37 | 2 | 0 | 0.001 | 3.279 | 0.278 |
| Glycerolipid metabolism | 34 | 0 | 3 | 0.044 | 1.353 | 0.242 |
| Glycine, serine and threonine metabolism | 72 | 0 | 4 | 0.089 | 1.052 | 0.225 |
| Nitrogen metabolism | 10 | 0 | 1 | 0.170 | 0.769 | 0.222 |
| Butanoate metabolism | 29 | 0 | 2 | 0.130 | 0.885 | 0.214 |
| Phenylalanine metabolism | 21 | 0 | 1 | 0.403 | 0.395 | 0.200 |
| Vitamin B6 metabolism | 21 | 0 | 1 | 0.430 | 0.366 | 0.200 |
| Cysteine and methionine metabolism | 72 | 0 | 6 | 0.007 | 2.185 | 0.197 |
| Valine, leucine and isoleucine biosynthesis | 12 | 0 | 1 | 0.170 | 0.769 | 0.182 |
| Drug metabolism - other enzymes | 69 | 0 | 4 | 0.096 | 1.019 | 0.162 |
| Tyrosine metabolism | 88 | 0 | 1 | 0.889 | 0.051 | 0.161 |
| Citrate cycle (TCA cycle) | 42 | 0 | 1 | 0.645 | 0.190 | 0.146 |
| Glycosaminoglycan biosynthesis - heparan sulfate or heparin | 15 | 0 | 1 | 0.279 | 0.554 | 0.143 |
| Glycosaminoglycan biosynthesis - chondroitin sulfate or dermatan sulfate | 18 | 0 | 1 | 0.312 | 0.506 | 0.118 |
| Arachidonic acid metabolism | 93 | 0 | 2 | 0.678 | 0.168 | 0.109 |
| Drug metabolism - cytochrome P450 | 39 | 0 | 1 | 0.571 | 0.243 | 0.105 |
| Steroid hormone biosynthesis | 177 | 0 | 2 | 0.950 | 0.022 | 0.102 |
| Nicotinate and nicotinamide metabolism | 43 | 0 | 1 | 0.734 | 0.134 | 0.095 |
| Retinol metabolism | 45 | 0 | 1 | 0.758 | 0.120 | 0.091 |
| Propanoate metabolism | 46 | 0 | 1 | 0.678 | 0.169 | 0.089 |
| Pyruvate metabolism | 48 | 0 | 1 | 0.693 | 0.159 | 0.085 |
| Galactose metabolism | 51 | 0 | 1 | 0.678 | 0.169 | 0.080 |
| Alanine, aspartate and glutamate metabolism | 61 | 0 | 1 | 0.791 | 0.102 | 0.067 |
| Histidine metabolism | 32 | 0 | 1 | 0.528 | 0.277 | 0.065 |

| | | | | | | |
|---|---|---|---|---|---|---|
| Arginine and proline metabolism | 74 | 0 | 3 | 0.247 | 0.608 | 0.055 |
| Steroid biosynthesis | 81 | 0 | 1 | 0.851 | 0.070 | 0.050 |
| beta-Alanine metabolism | 44 | 0 | 1 | 0.662 | 0.179 | 0.047 |
| Fatty acid biosynthesis | 129 | 0 | 1 | 0.983 | 0.008 | 0.047 |
| Fatty acid degradation | 102 | 0 | 1 | 0.952 | 0.021 | 0.040 |
| Glycosphingolipid biosynthesis - lacto and neolacto series | 104 | 0 | 2 | 0.774 | 0.111 | 0.039 |
| Porphyrin metabolism | 55 | 0 | 1 | 0.678 | 0.169 | 0.037 |
| Lysine degradation | 57 | 0 | 1 | 0.721 | 0.142 | 0.036 |
| Metabolism of xenobiotics by cytochrome P450 | 118 | 0 | 2 | 0.720 | 0.143 | 0.034 |
| Folate biosynthesis | 60 | 0 | 1 | 0.791 | 0.102 | 0.034 |
| One carbon pool by folate | 72 | 0 | 1 | 0.889 | 0.051 | 0.028 |
| Tryptophan metabolism | 84 | 0 | 1 | 0.871 | 0.060 | 0.024 |
| Glycerophospholipid metabolism | 87 | 0 | 1 | 0.913 | 0.039 | 0.023 |

**Supplementary Table 11. Integrative Analysis of Significant Genes & Metabolites *mdx* RIB/VEH.**

| Pathway | Total metabolites & genes in pathway | Metabolite hits | Gene hits | Raw *p* value | -log10(*p*) | Impact |
|---|---|---|---|---|---|---|
| **Pathways with Gene and Metabolite Hits and Impact Score >0.2** | | | | | | |
| Galactose metabolism | 51 | 2 | 2 | 0.057 | 1.238 | 0.84 |
| Fatty acid degradation | 102 | 1 | 3 | 0.704 | 0.152 | 0.316 |
| Butanoate metabolism | 29 | 1 | 2 | 0.089 | 1.048 | 0.25 |
| Pentose and glucuronate interconversions | 34 | 1 | 2 | 0.111 | 0.953 | 0.363 |
| Citrate cycle (TCA cycle) | 42 | 1 | 2 | 0.219 | 0.658 | 0.292 |
| Propanoate metabolism | 46 | 1 | 2 | 0.257 | 0.589 | 0.222 |
| **Pathways with Gene and Metabolite Hits and Impact Score <0.2** | | | | | | |
| Starch and sucrose metabolism | 37 | 1 | 1 | 0.508 | 0.293 | 0.194 |
| Taurine and hypotaurine metabolism | 17 | 1 | 1 | 0.142 | 0.845 | 0.187 |
| Steroid hormone biosynthesis | 177 | 1 | 4 | 0.870 | 0.060 | 0.153 |
| Arginine biosynthesis | 28 | 1 | 1 | 0.260 | 0.584 | 0.148 |
| Alanine, aspartate and glutamate metabolism | 61 | 1 | 2 | 0.433 | 0.363 | 0.083 |
| **Other Pathways** | | | | | | |
| Fatty acid biosynthesis | 129 | 0 | 2 | 0.988 | 0.005 | 1.039 |
| Neomycin, kanamycin and gentamicin biosynthesis | 4 | 0 | 1 | 0.139 | 0.853 | 0.666 |
| Metabolism of xenobiotics by cytochrome P450 | 118 | 0 | 5 | 0.353 | 0.451 | 0.598 |
| Ascorbate and aldarate metabolism | 19 | 0 | 2 | 0.134 | 0.872 | 0.444 |
| Linoleic acid metabolism | 17 | 0 | 3 | 0.050 | 1.300 | 0.375 |
| Glycosaminoglycan degradation | 44 | 0 | 4 | 0.059 | 1.226 | 0.372 |
| Pyruvate metabolism | 48 | 0 | 4 | 0.100 | 0.996 | 0.297 |
| Sphingolipid metabolism | 81 | 0 | 5 | 0.279 | 0.554 | 0.275 |
| Mucin type O-glycan biosynthesis | 27 | 0 | 3 | 0.061 | 1.208 | 0.269 |
| Drug metabolism - other enzymes | 69 | 0 | 5 | 0.145 | 0.836 | 0.25 |
| Glycerolipid metabolism | 34 | 0 | 3 | 0.136 | 0.865 | 0.242 |
| Glycolysis or Gluconeogenesis | 60 | 0 | 4 | 0.228 | 0.640 | 0.237 |
| Retinol metabolism | 45 | 0 | 4 | 0.167 | 0.777 | 0.227 |
| Nitrogen metabolism | 10 | 0 | 2 | 0.028 | 1.547 | 0.222 |
| Arachidonic acid metabolism | 93 | 0 | 7 | 0.062 | 1.202 | 0.217 |
| Glycosphingolipid biosynthesis - ganglio series | 47 | 0 | 2 | 0.575 | 0.239 | 0.217 |
| Ether lipid metabolism | 40 | 0 | 4 | 0.0501 | 1.294 | 0.205 |
| Cysteine and methionine metabolism | 72 | 0 | 6 | 0.056 | 1.248 | 0.197 |
| Drug metabolism - cytochrome P450 | 39 | 0 | 4 | 0.035 | 1.446 | 0.184 |
| Valine, leucine and isoleucine biosynthesis | 12 | 0 | 1 | 0.260 | 0.583 | 0.181 |
| One carbon pool by folate | 72 | 0 | 2 | 0.863 | 0.063 | 0.169 |
| Inositol phosphate metabolism | 69 | 0 | 4 | 0.312 | 0.505 | 0.161 |
| Glutathione metabolism | 57 | 0 | 3 | 0.352 | 0.452 | 0.160 |

| | | | | | | |
|---|---|---|---|---|---|---|
| Amino sugar and nucleotide sugar metabolism | 93 | 0 | 2 | 0.899 | 0.045 | 0.152 |
| Glycosaminoglycan biosynthesis - heparan sulfate or heparin | 15 | 0 | 1 | 0.410 | 0.386 | 0.142 |
| Glycine, serine and threonine metabolism | 72 | 0 | 2 | 0.779 | 0.108 | 0.140 |
| Glycosaminoglycan biosynthesis - chondroitin sulfate or dermatan sulfate | 18 | 0 | 1 | 0.454 | 0.342 | 0.117 |
| Glycerophospholipid metabolism | 87 | 0 | 4 | 0.515 | 0.287 | 0.116 |
| Glycosphingolipid biosynthesis - globo and isoglobo series | 37 | 0 | 2 | 0.407 | 0.390 | 0.111 |
| Fructose and mannose metabolism | 37 | 0 | 1 | 0.764 | 0.116 | 0.111 |
| alpha-Linolenic acid metabolism | 22 | 0 | 2 | 0.134 | 0.872 | 0.095 |
| Purine metabolism | 173 | 0 | 3 | 0.985 | 0.006 | 0.093 |
| Pentose phosphate pathway | 49 | 1 | 0 | 0.153 | 0.815 | 0.083 |
| Steroid biosynthesis | 81 | 0 | 2 | 0.803 | 0.094 | 0.075 |
| Porphyrin metabolism | 55 | 0 | 2 | 0.531 | 0.274 | 0.074 |
| Glyoxylate and dicarboxylate metabolism | 56 | 0 | 1 | 0.839 | 0.075 | 0.072 |
| Pantothenate and CoA biosynthesis | 36 | 1 | 0 | 0.134 | 0.871 | 0.057 |
| Pyrimidine metabolism | 100 | 1 | 0 | 0.246 | 0.607 | 0.040 |
| Fatty acid elongation | 75 | 0 | 1 | 0.950 | 0.022 | 0.040 |
| Arginine and proline metabolism | 74 | 0 | 2 | 0.779 | 0.108 | 0.027 |
| N-Glycan biosynthesis | 81 | 0 | 1 | 0.957 | 0.018 | 0.025 |
| Tryptophan metabolism | 84 | 0 | 1 | 0.963 | 0.016 | 0.024 |
| Tyrosine metabolism | 88 | 0 | 1 | 0.971 | 0.012 | 0.022 |
| Glycosphingolipid biosynthesis - lacto and neolacto series | 104 | 0 | 1 | 0.990 | 0.004 | 0.019 |
| Primary bile acid biosynthesis | 90 | 1 | 0 | 0.284 | 0.545 | 0.011 |
